# EphB3 receptor negatively regulates osteogenesis in mice

**DOI:** 10.1101/2023.06.15.544777

**Authors:** Mariano R. Rodríguez-Sosa, David Alfaro, Luis M. del Castillo, Adrián Belarra, Agustín G. Zapata

## Abstract

Bone homeostasis is a complex process in which some Eph kinase receptors and their Ephrin ligands appear to be involved. In the present study, we address this issue by examining the capacity of adipose tissue-derived mesenchymal stromal cells (Ad-MSC) derived from either WT, *EphB2*-and *EphB3*-KO mice to differentiate into bone tissue. Differentiation capacities were evaluated in cultured MSC by RT-qPCR and histological staining, revealing that whereas *EphB2*^-/-^ MSC cultured in a specific medium expressed mainly pro-adipogenic transcription factors, *EphB3*^-/-^ MSC showed abundant osteogenic transcripts, such as *Runx2*, *Msx2* and *Osterix.* In addition, the lack of EphB3 signaling alters the genetic profile of differentiating Ad-MSC, reducing the expression of many inhibitory molecules and antagonists of the BMP signaling pathway, and increasing *Bmp7* expression, a robust bone inductor. Then, to confirm the osteogenic role of EphB3 *in vivo*, we studied the condition of two animal models of induced osteoporosis (ovariectomy or long-term glucocorticoid treatment). Interestingly, in both models, both WT and *EphB2*^-/-^ mice equally developed the disease but *EphB3*^-/-^ mice did not exhibit the typical bone loss, nor did they show increased urine Ca^2+^ or blood serum CTX-1. The proportions of osteoprogenitor cells and pre-osteoblasts were also found to be significantly higher in *EphB3*-KO mice, and the osteoclasts significantly reduced, as compared to WT and *EphB2*-KO mice. We conclude that EphB3 acts as a negative regulator of the osteogenic differentiation, and its absence prevents the development of experimentally-induced osteoporosis.

## Introduction

Mesenchymal stromal/stem cells (MSC) are a heterogeneous cell population of stromal cells capable of plastic adherence, under standard culture conditions, and multilineage cell differentiation mainly into osteogenic, adipogenic and chondrogenic cells. They can be isolated from almost any organ containing connective tissue [1], and are phenotypically characterized by the expression of non-exclusive cell markers and the lack of others [2]. Due to their remarkable immunomodulatory properties and capacities to give rise to mesoderm-derived tissues, MSC are currently an excellent therapeutic tool [3-5].

Numerous morphogenetic factors (i.e., BMP, Wnt/β−catenin, Igf-1, etc.) are involved in the differentiation of MSC to bone cell lineage, with BMP signaling being one of the main routes involved in this process [6]. Other molecules implicated in osteogenic differentiation are the Eph (Erythropoietin-producing hepatocellular carcinoma) receptors and their ligands, Ephrins [7-12]. Eph constitute the largest family of tyrosine kinase receptors in animal cells. They are classified into two subfamilies (A and B) that, together with their ligands, Ephrins, are involved in numerous processes during embryonic development and in adult tissue homeostasis [13, 14]. Eph and Ephrins can signal bidirectionally, which means that both receptors and ligands are able to simultaneously send intracellular signals through Eph (forward signal) and Ephrin (reverse signal) [15, 16], regulating cell attraction/repulsion, cell adhesion, survival, cell proliferation and differentiation [17]. Both MSC and bone cells express Eph and Ephrins A and B and their lack results in altered bone development and malformations [18-21]. However, their specific functions in osteogenesis are not conclusively known [7]. Both EphA2 [8, 12, 22] and EphA5 [23] have been reported as negative regulators of the osteogenic differentiation of MSC, whereas global elimination of EphA4 results in reduced epiphyseal growth plates [7], and bidirectional signaling of the EphA2/EphrinA2 pair regulates bone remodeling [22]. With regards to the B family, it has been proposed [8] that forward signaling via EphB4, expressed on osteoblasts, and reverse signals through EphrinB2, expressed in osteoclasts, regulate bone formation and reabsorption [9, 12]. EphB2 and EphrinB1 were also involved, although their participation appears to be less relevant [24]. On the other hand, treatment with EphB2-Fc fusion proteins increased osteogenesis, whereas blocking EphB1 or EphB4 signaling peptides inhibited it [24].

On the other hand, information about a possible role for EphB3 in osteogenesis and bone homeostasis is scarce. Together with EphB2, a role for EphB3 in the fusion of the secondary palate was pointed out [25, 26]. More recently, strong EphB3 expression in adult calvaria sutures and the proliferative chondrocytes of long bones has been shown [27]. These authors have reported that EphB3 might limit osteogenesis because its lack results in increased bone volume [27], but they also remarked that the effect is “relatively minimal and presumably transient”.

In the present study, we comparatively analyze the role of EphB2 and EphB3 in the differentiation of adipose tissue-derived MSC (Ad-MSC) into bone cell lineage. Our results confirm that the EphB3 signaling pathway is a potent suppressor of osteogenic differentiation, because its lack produces important augmented osteogenic activity, by inducing decreased expression of several BMP antagonists and negative regulators, and an increase in *Bmp7*. This behavior prevents the development of experimentally-induced osteoporosis in *EphB3*^-/-^ mice, by increasing the proportions of osteogenic progenitor cells and reducing those of osteoclasts. These results point to EphB3 signaling as a possible new target for the treatment of bone deficits, including fractures and osteoporosis: important public health problems that affect 200 million people worldwide [28].

## Materials and methods

### 1) Animals and ethical procedures

Only healthy WT, *EphB2*-(*EphB2*^-/-^) or *EphB3*-deficient (*EphB3*^-/-^) mice (*Mus musculus*, L. 1758), with a CD1 genetic background (outbreed) and weighing around 30 g, were used in the present study. All KO used animals were previously genotyped by PCR. The original animals were kindly donated by Dr. Henkemeyer (*Southwestern Medical Center*, Dallas, TX, USA) and are currently part of our own colony.

Animals were housed under sterile conditions in ventilated cages at the Animal Housing facilities of the UCM, with a maximum caging density of five adult mice from the same litter and sex, environmental enrichment (nesting material, tubular shelter and social housing), sterile *ad libitum* food (hard pellets) and water, controlled temperature and humidity (24 °C, 55%) and 12/12h light/dark cycle, according to current animal care policies. Excluding animal criteria and humane endpoints were established before the experiment starts and were mainly related to dramatic body-weight loss, important behavioral impairments or if the animal died prematurely, preventing the collection of data. In any case, no animals were excluded in this study. All procedures used here were approved by the Ethics Committee for Animal Research of the UCM and Regional Government of Madrid. This study is in accordance with the ARRIVE guidelines 2.0.

### 2) Isolation and culture of MSC derived from murine adipose tissue

For every Ad-MSC isolation, we used a mixed pool of five 4-6-month-old WT or *EphB* mutant mice of both sexes. This protocol was carried out 5 times, using 25 mice per group and 75 in total. Mice were sacrificed by cervical dislocation and adipose tissue from the abdominal zone was isolated. After physical disaggregation and digestion with 2 mg/ml collagenase B (*Roche*, CA, USA), the enzymatic activity was stopped by diluting with PBS 1X, cell suspension was filtered using a 70 μm filter (*BD-Bioscience*, NJ, USA) into a 50 ml Falcon tube (*BD-Bioscience*) and centrifuged for 5 min, at 1500 rpm and 10 °C. Then, the supernatant was removed, the pellet resuspended in culture medium, plated on T150 (Corning, AZ, USA) with *Dulbecco’s Modified Eagle Medium* (DMEM) (*Hyclone*, UT, USA) supplemented with 10% fetal bovine serum (FBS) (Gibco, MA, USA), 1X pyruvate (Sigma-Aldrich, Germany), 1X antibiotic-antimycotic (Lonza, Switzerland) and 1X L-glutamine (Lonza), and cultured at 37 °C and 5% CO2. After 24h the flasks were washed twice with PBS 1X (Lonza) and the culture medium was replaced by supplemented *Mesencult medium* (*Stemcell*, France) according to the manufacturer’s instructions. Cells were cultured until 80-90% confluency, detached using trypsin 0.025% (*Hyclone*) and seeded in *Mesencult medium* (*Stemcell*) for another passage. On each new passage, Ad-MSC were kept in DMEM (*Hyclone*) supplemented media as described before, changing the medium every 5-6 days or until reaching cell confluency. The growth of the cells was monitored using an inverted phase contrast microscope *Nikon Eclipse TS100* (*Nikon*, Japan). For all the experiments, cells in passage 3 or 4 were used. According to the Reduction principle (3Rs), the excess of cells was frozen for use in other related studies.

### 3) Growth curve and doubling time

We seeded 10000 Ad-MSC per well in a P24 plate (*Corning*) with DMEM medium (*Hyclone*) supplemented with 10% fetal bovine serum (FBS) (*Gibco*), 1X pyruvate (*Sigma-Aldrich*), 1X antibiotic-antimycotic (*Lonza*) and 1X L-glutamine (*Lonza*) and cultured at 37 °C and 5% CO2. On days 4, 8, 12 and 21, cells were detached with trypsin (*Hyclone*), centrifuged at 1500 rpm for 5 min and resuspended in DMEM to be counted in a *Neubauer* chamber, using trypan blue to exclude dead cells. To determine the doubling time, we counted live cells on each passage using the following equation:

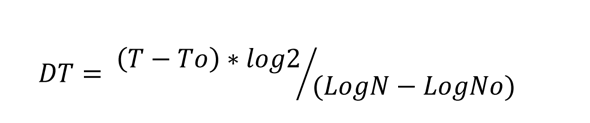

Where “DT” is doubling time, “T” and “To” is culture time in hours, “N” is number of cells after incubation and “No” the initial seeded cells [29].

### 4) Osteoblast isolation

We used a modified protocol described by Baryanwo *et al.* (2019) [30]. Briefly, left femurs from 4–6-month-old female control or dexamethasone-induced osteoporotic mice (see further) were harvested. The bone marrow was flushed using a 25G syringe and reserved for osteoclast analysis. Then, the bone was cut into small pieces and digested with 2 mg/ml *Stemzyme* (*Worthington*, NJ, USA) for 30 min at 37 °C. Cell suspensions were then filtered using a 70 µm filter (*BD-Bioscience*) and centrifuged for 5 min at 1500 rpm and 10 °C. After centrifugation, cells were resuspended on an erythrocyte lysis buffer for 5 min at 4 °C, centrifuged again, resuspended in DMEM with 2% FBS for cell counting using a hemocytometer, and stained for flow cytometry analysis.

### 5) Osteoclast isolation and Tartrate-Resistant Acid Phosphatase (TRAP) staining

For osteoclast isolation we used a modified protocol described by Tevlin *et al.* (2014) [31]. Briefly, a density gradient cell separation of the reserved flushed-bone marrows from osteoblast isolations was performed using *Histopaque-1077* (*Sigma-Aldrich*) for 15 min at 1500 rpm and room temperature (RT). The cells from the buffy coat were removed and centrifuged again at 1500 rpm and 4°C for 5 min, setting 2×10^6^ cells aside for flow cytometry staining. Bone marrow cells were seeded in 48-well plates (*Corning*) at a density of 0.15×10^6^ cell/well and cultured for 2 days in supplemented α-MEM with 30 ng/ml M-CSF (*PeproTech*, UK). Media were replaced with α-MEM supplemented with 30 ng/ml M-CSF and 30 ng/ml RANKL (*PeproTech*) for 3 days. Media were replaced again with α-MEM supplemented simultaneously with 30 ng/ml M-CSF, 30 ng/ml RANKL and 10 ng/ml TNF-α (*PeproTech*) for 2 days more. After 7 days of culture, cells were stained with the TRAP-staining kit (*Sigma-Aldrich*) following the manufactureŕs instructions. Osteoclasts were identified as TRAP^+^ cells with more than 3 nuclei, using an inverted phase contrast microscope *Nikon Eclipse TS100* (*Nikon*) with a *Motic3+* digital camera attached (*Motic*, Hong Kong) to take pictures and manually count osteoclasts.

### 6) Flow cytometry

Ad-MSC, isolated as described before, were characterized using a panel of antibodies (**Table 1**). Cell surface staining was performed using the primary and secondary antibodies included in the table, following supplier’s instructions. Both stainings were carried out for 15 min at 4 °C.

To determine the cell cycle, Ad-MSC were incubated for 30 min with 1/3 dilution of *Propidium Iodide* (PI) solution (*Cytognos*, Spain) in PBS 1X. For the assessment of cell viability, cells were incubated with diluted 7AAD (Sigma-Aldrich) (1/2000) and *Annexin-V* 634 (*Immunostep*, Spain).

Osteoblasts were phenotyped using the following cell surface markers: Sca1-FITC, CD51-PE, Lin-APC, CD45-647, CD31-APC (*Biolegend*, CA, USA). 1×10^6^ cells were stained for 15 min at 4 °C. Osteoclasts were identified by CD11b-FITC, cKit-PE, CD3-PerCP, CD45R-PerCP and CD115-APC (*Biolegend*) expression, using 2×10^6^ cells for staining for 15 min at 4 °C.

Flow cytometric analyses were performed using a *FACSCalibur* device (SCR_000401, *BD Biosciences*, CA, USA) equipped with *CellQuest* software at the Cytometry and Fluorescence Microscopy Centre (UCM).

### 7) Osteogenic and adipogenic differentiation of Ad-MSC

To induce osteogenic and adipogenic differentiation, Ad-MSC were cultured in specific *Mesencult differentiation media* (*Stemcell*). The differentiation was evaluated at 7 days (by RT-qPCR) and 21 days (by histological staining), changing the culture media every 3-5 days. Also, to further dissect the underlying signaling pathways involved in the osteogenic differentiation, we evaluated after 7 days of culture by RT-qPCR the transcript expression of several genes implicated in this process (* marked in Table 2).

### 8) Quantitative real-time polymerase chain reaction (RT-qPCR)

RNA was isolated from Ad-MSC using a *Total RNA Extraction Kit* (*Biotools*, Spain) according to the supplieŕs instructions. Total cDNA from these cells was transcribed with a *High-Capacity cDNA Reverse Transcription Kit* (*Applied Biosystems*, MA, USA) and RT-qPCR were performed using the specific primers indicated in Table 2, in a *QuantStudio 12 K FLEX* PCR system (*Applied Biosystems*) with *QuantStudio Real-Time PCR Software* v1.2.2., at the Genomic Centre of the UCM. The relative expression of each sample was normalized to ΔCT values and represented as 2^−ΔΔCt^ referred to their respective DMEM cultured Ad-MSC values.

### 9) Histological staining with Alizarin Red and Oil Red

For *Alizarin Red* staining, cells were washed with PBS 1X and fixed with 70% ethanol (*Sigma-Aldrich*) for 1 h at 4 °C. Then, *Alizarin Red* (*Sigma-Aldrich*) work solution was added and incubated for 20 min at RT and protected from light. Finally, cells were washed with distilled water and pictures were taken with a Motic3+ digital camera (*Motic*) attached to an inverted microscope. To quantify the *Alizarin Red*, a 10% cetylpyridinium chloride (CPC) solution (*Sigma-Aldrich*) was added and shaken for 30 min at RT and protected from light. Then, 200 μl of this solution were transferred to a 96-well ELISA plate (*Corning*) to read absorbance at 562 nm on a *Biochrom Asys UVM 340* microplate reader (*Biochrom*, UK). The obtained values were compared with a standard curve to assess the concentration of *Alizarin* red for each assay.

For *Oil Red* staining, cells in each well were fixed with 10% formaldehyde (*Sigma-Aldrich*) for 20 min at RT and 70% isopropanol (*Sigma-Aldrich*) was added for 5 min. All reagents were removed, and *Oil Red* (*Sigma-Aldrich*) work solution was added and incubated for 5 min at 4 °C. Finally, the *Oil Red* solution was removed, and each well was washed with PBS 1X to take pictures as described before.

### 10) Experimental induction of osteoporosis

Both 3–4-month-old female WT and *EphB* KO mice were individually analyzed in this assay. Every group consisted of 6 animals, randomly assigned with a computer-based random order generator, using 54 mice in total. In any case, no adverse events occur in these mice.

For the ovariectomy-induced osteoporosis, mice were divided into a control group and an ovariectomized (OVX) group. Mice were anesthetized by intraperitoneal injection of a ketamine–xylazine solution (*Ketamine*: *Ketolar* 50 mg/mL, *Pfizer Group*, Spain; *Xylazine*: *Rompun* 2%, *Bayer*, Germany). Animals were put away from the light while anesthesia. After this, bilateral ovariectomy was performed using two incisions in the mouse’s back that were closed using suture. For the control group, just the incisions were made but the ovaries were not removed. To reduce pain after surgery, analgesics (Acetaminophen 1mg/ml) were provided during the first week diluted in sterile water for voluntary ingestion. Two months later, mice were sacrificed.

For the dexamethasone induced-osteoporosis, mice were divided into two groups: a control and a dexamethasone (*Calier*, Spain) treated group (DEX). 2.5 mg/kg/day of corticoid was injected intraperitoneally using a 27G needle in a 1 ml syringe daily for 14 days. The control group was injected with PBS 1X. Mice were sacrificed at day 15.

In both models, the right femur and the second lumbar vertebrae were extracted for micro-CT analysis, and urine and blood were collected for other studies. Because the results obtained for both osteoporotic mouse models were similar, after initial analyses, only DEX treated mice were evaluated. After verifying the osteoporotic phenotype by micro-CT, blood serum, urine and left femurs were used to obtain osteoblast and osteoclast suspensions, from WT and EphB-KO mice. Both WT and *EphB* KO 3–4-month-old female mice were individually used in this assay. Every analyzed group was formed by 5 animals, using 30 mice in total.

The daily monitorization of the animals after surgery and during the dexamethasone injection was performed by the research team and the staff from the Animal Housing facility.

We chose these two osteoporotic mouse models because in humans this pathology is largely related to two main findings: reduced estrogen levels (in menopause women) or long-term corticoid treatments for immunosuppression.

### 11) Sample collection and preparation

Collected blood samples were allowed to clot to obtain sera by centrifugation at 3000 rpm for 5 min. Serum and urine samples were stored at −20 °C for further analysis.

### 12) Enzyme linked immunosorbent assay (ELISA)

*RatLaps EIA* kits were used to determine serum carboxy-terminal telopeptide of type I collagen (CTX-1) (*Immunodiagnostics Systems*, UK), calcium (*Abcam*, UK) and estrogens (*Abcam*) following manufactureŕs instructions (*Immunodiagnostics Systems*, UK) and read at the appropriate wavelength using a *Biochrom Asys UVM 340* microplate reader (*Biochrom*, UK).

### 13) Micro-CT analysis

Micro-CT analysis were blinded during data collection and quantification. Tomographic acquisitions of formaldehyde-fixed samples were visualized in a micro-CT system consisting of a *Hamamatsu* brand X-ray tube with a focus size of 20 μm and a flat-panel with a pixel size of 50 μm. The acquisition of 2D-projections was set at 50 kV and 100 μA for an exposure time of 1200 ms through a 360°-turn (0.36°/step). The 3D image stacks with a voxel size of 12.5 μm were reconstructed using *Octopus Reconstruction software* (*TESCAN XRE*, Belgium) which allows flat-field, ring artifact corrections and segmentation into binary images (8-bit BMP images) for the subsequent image processing. Quantification of bone volume fraction (BV/TV) and trabecular thickness (Tb.th) was performed for each vertebra and femur, selecting at least 50 slices from each bone stack with *BoneJ* (SCR_018166, *ImageJ plug-in* [32]), and the 3D images of every scanned bone was obtained with *Dragonfly software* (CA, USA).

### 14) Statistical analysis

Animal sample sizes were calculated *a priori* according to the established formula for analysis of variance. Statistical interpretation of the data was performed by one-way or two-way analysis of variance (ANOVA) with a subsequent *Tukey test* as a post-test, depending on the experiment, after evaluating normality by *Shapiro Wilk test*. Data are represented as means ± standard deviations. Statistically significant differences were set for a *p value* less than 0.05, showing asterisks (*) when *EphB*-deficient and control WT values were compared, and triangles (Δ) for *EphB2* vs *EphB3* mutant values. All calculations were performed using *GraphPad software* (SCR_002798, *GraphPad Prism*, CA, USA).

### 15) Cell cartoon credits (Fig. 10 and 11)

Cartoons were elaborated by *Laboratoires Servier-Smart Servier* website: Images related to Osteoblasts and Osteoclast progenitors, Bone structure and Bones. CC BY-SA 3.0. https://commons.wikimedia.org/w/index.php?curid=82640795 and …82640809.

## Results

### Characterization of Ad-MSC of both WT and EphB-KO mice

Ad-MSC of either WT, *EphB2*^-/-^ or *EphB3*^-/-^ mice were cultured *in vitro* and analyzed after the second passage. All the cultured cells had the expected fibroblastic morphology **(Fig. 1a**) and similar phenotypic profiles, as evaluated by flow cytometry, expressing CD29, CD44, CD73, CD105, CD106 and Sca1, but not CD11b and CD45 (**Fig. 1b**).

**Fig. 1.**
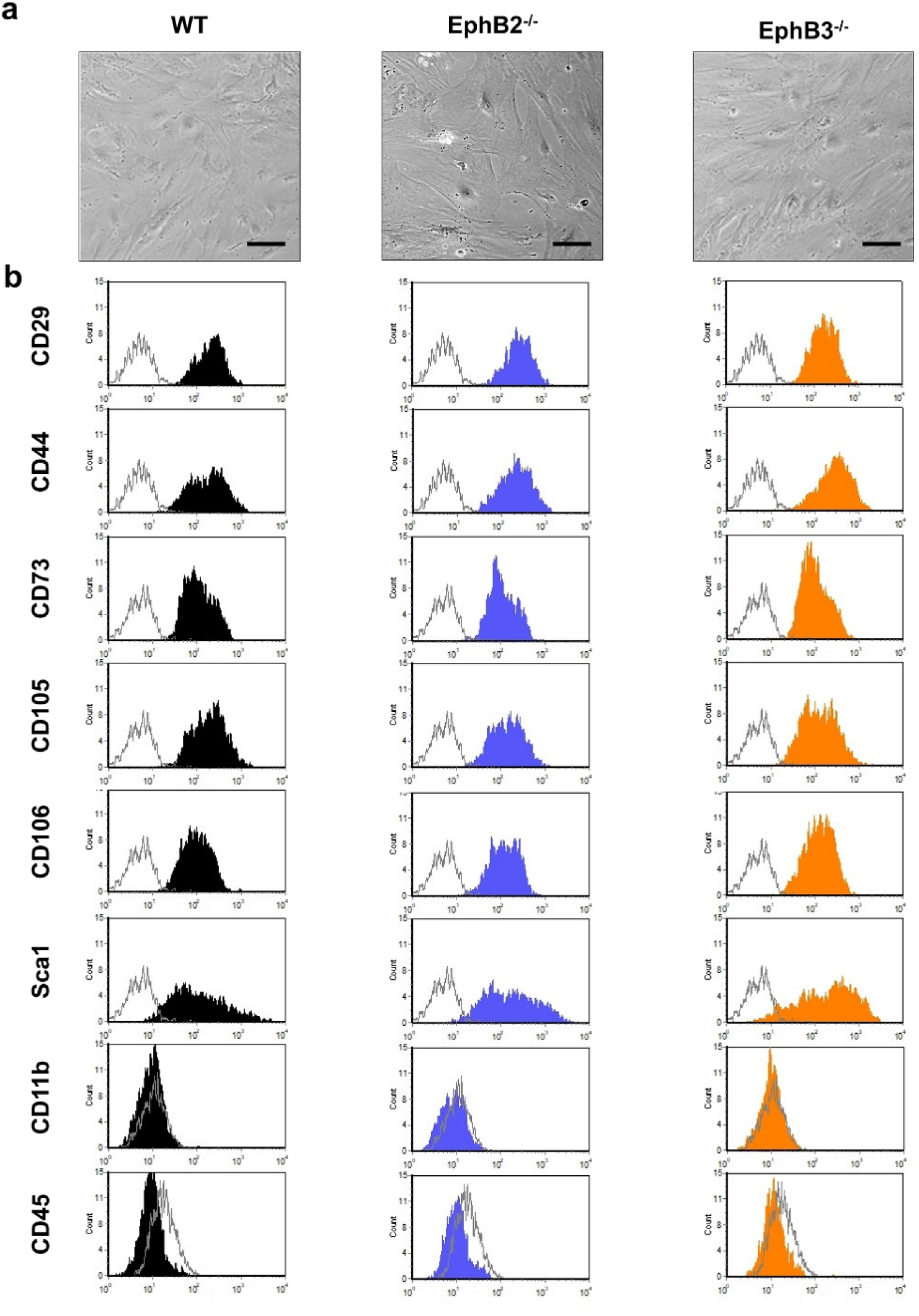
Morphology and cell surface marker expression of Ad-MSC derived from WT, *EphB2*^-/-^ and *EphB3*^-/-^ mice. No significant changes in cell morphology or in the expression profile of MSC markers were found between *EphB*-deficient and WT mice. (a) *In vitro* morphology of Ad-MSC near cell confluency (100X, Scale: 200 μm). (b) Representative histograms of the cell surface marker expression profiles of WT and mutant Ad-MSC is shown. The colored regions represent the expression levels of each cell marker and empty histograms are the negative controls (n=5, with 5 mice per group).

**Table 1.** Cell surface antibodies used for flow cytometry analysis.

| Antibody | Conjugation | Supplier | Cell staining | RRID |
| --- | --- | --- | --- | --- |
| **CD11b** | FITC | BioLegend | MSC / Osteoclast | AB_312789 |
| **CD44** | Alexa Fluor® 488 |  | MSC | AB_493679 |
| **CD45** | Alexa Fluor® 647 |  | MSC / Osteoblast | AB_493534 |
| **Sca1** | FITC |  | MSC / Osteoblast | AB_313342 |
| **CD29** | IgG<sub>2A</sub> | R&D Systems | MSC | AB_312878 |
| **CD73** | IgG<sub>2A</sub> |  |  | AB_2904273 |
| **CD105** | IgG<sub>2A</sub> |  |  | AB_961066 |
| **CD106** | IgG<sub>2A</sub> |  |  | AB_313202 |
| **Goat anti-rat IgG** | Alexa Fluor® 488 | Molecular Probes |  | AB_141373 |
| **CD3** | PerCP | BioLegend | Osteoclast | AB_893319 |
| **CD31** | APC |  | Osteoblast | AB_312905 |
| **CD45R** | PerCP |  | Osteoclast | AB_893353 |
| **CD51** | PE |  | Osteoblast | AB_2129493 |
| **CD115** | APC |  | Osteoclast | AB_2085222 |
| **cKIT** | PE |  | Osteoclast | AB_313216 |
| **Lineage** | APC |  | Osteoblast | AB_1645213 |

**Table 2.**
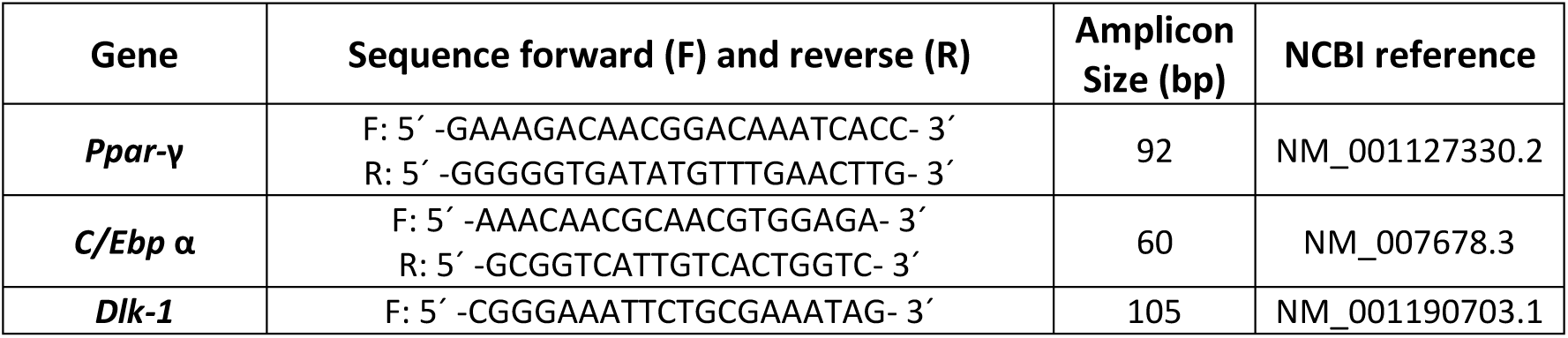
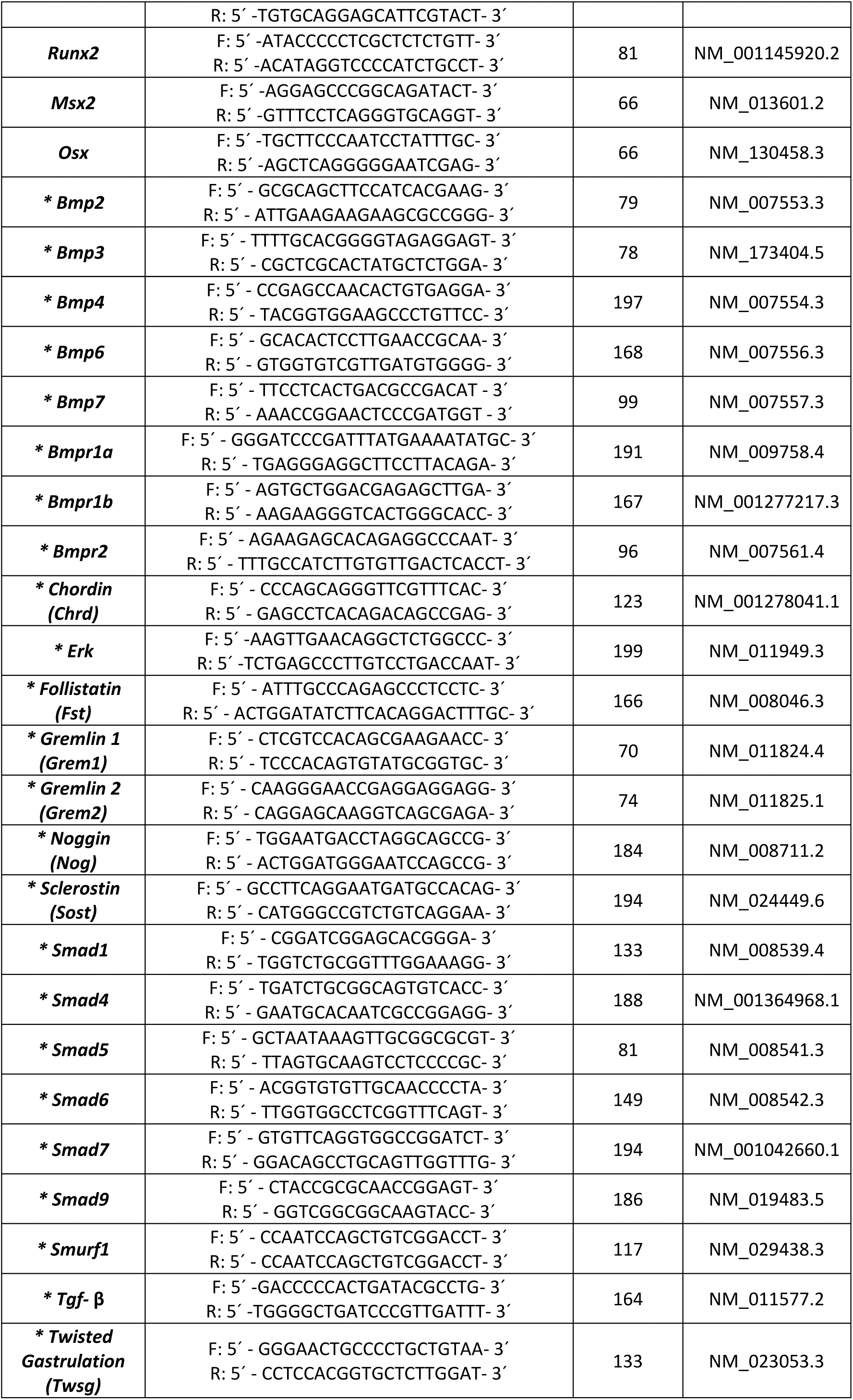
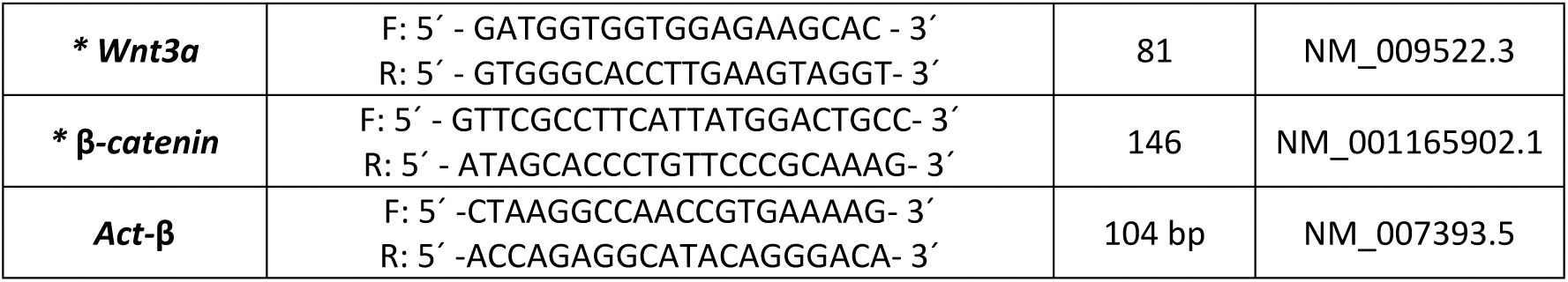
Primer sequences used for molecular characterization by RT-qPCR of genes involved in the differentiation of Ad-MSC to either adipogenic or osteogenic cell lineages, including those related to additional signaling pathways (* marked).

We also analyzed the growth curve of Ad-MSC for 21 days, until full confluency was achieved in a P24-well plate. No changes were found between *EphB*-deficient and WT Ad-MSC in either growth (**Fig. 2a**) and doubling-time (**Fig. 2b**), or in the percentage of dividing (**Fig. 2c**) and apoptotic cells (**Fig. 2d, 2e**).

**Fig. 2.**
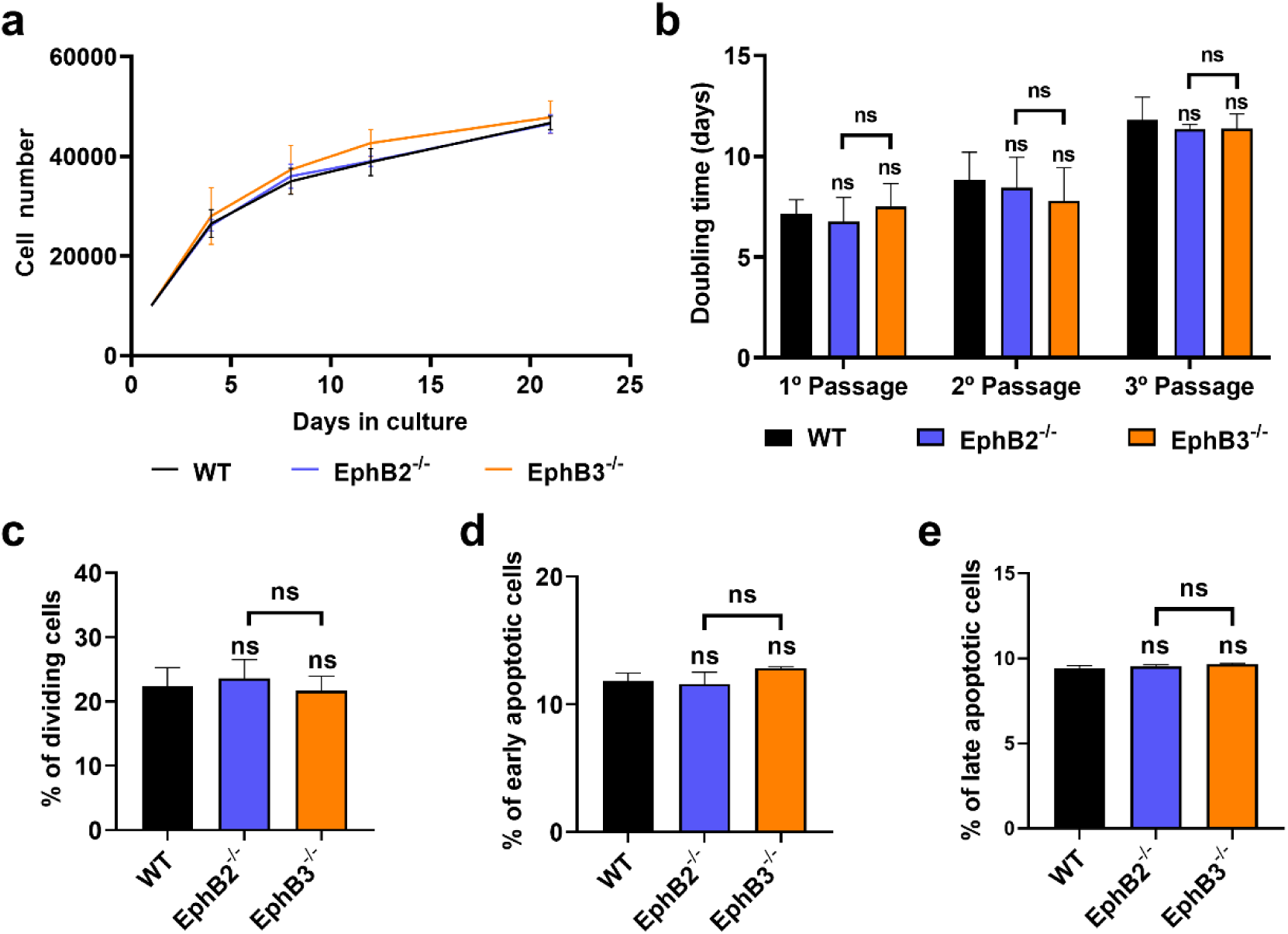
The absence of EphB2 or EphB3 does not modify the cell homeostasis of Ad-MSC, as demonstrates the analysis of growth, doubling time, cell proliferation or survival. (a) Growth curve of cultured cells at different days. (b) Doubling time of the studied Ad-MSC at different passages. (c) Percentages of cycling Ad-MSC of different genotypes. (d, e) Proportions of early (Annexin^+^/7AAD^-^) (d) and late (Annexin^+^/7AAD^+^) (e) apoptotic Ad-MSC in WT and *EphB*-KO cultures at passage 3-4 (ns=no significant difference) (n=5, with 5 mice per group).

### EphB2^-/-^ and EphB3^-/-^ Ad-MSC show different in vitro capacities to differentiate into bone and adipose tissue

First, we analyzed by RT-qPCR the expression of transcription factors known to be related to adipogenic differentiation in Ad-MSC cultured for 7 days in adipogenic medium. Pro-adipogenic factors (*Ppar*-γ and *C/Ebp*-α) and the repressor gene *Dlk-1* were studied. *EphB2*^-/-^ Ad-MSC had a significantly higher expression of *Ppar*-γ and *C/Ebp*-α and reduced values of *Dlk-1*, than *EphB3*^-/-^ Ad-MSC (**Fig. 3a-c**). No differences were found between WT and *EphB3*^-/-^ Ad-MSC in the expression of these pro-adipogenic factors **(Fig. 3a, 3b)**, but *EphB3* mutant cells showed a significantly higher expression of the repressor gene *Dlk-1* than the WT cells **(Fig. 3c).**

**Fig. 3.**
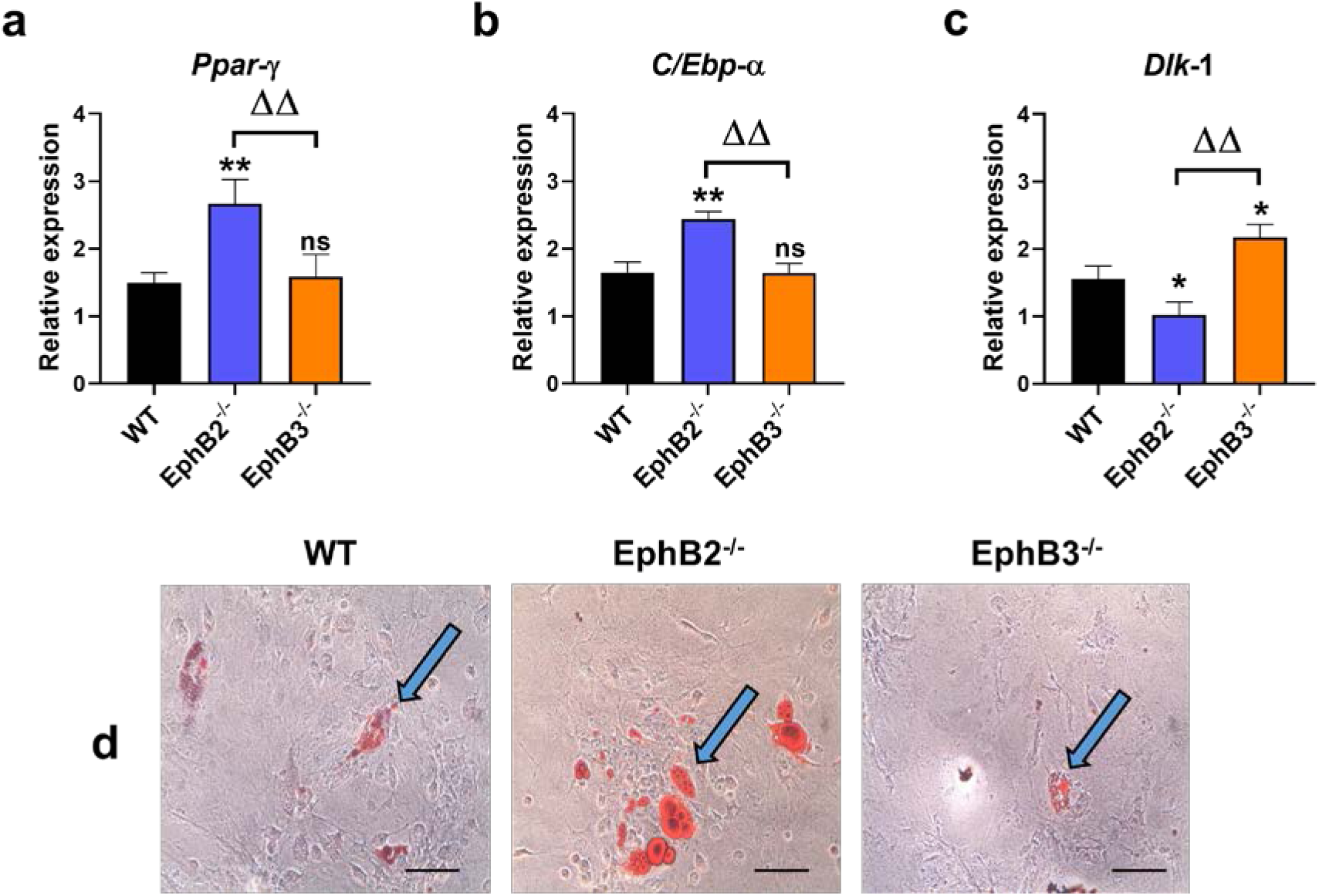
Adipogenic differentiation of both WT and *EphB*-deficient Ad-MSC. (a, b, c) Expression of adipogenic transcription factors in WT and *EphB*-KO Ad-MSC after 7 days cultured in adipogenic medium. A higher expression of the pro-adipogenic *Ppar*-γ and *C/Ebp*-α transcripts and a lower expression of the *Dlk-1* repressor were observed in EphB2^-/-^ Ad-MSC. On the contrary, *EphB3*^-/-^ Ad-MSC increased the expression of the inhibitor *Dlk-1* during the adipogenic differentiation (ns=no significant difference, */Δ p value<0.05). (d) Cultures were stained with *Oil Red* to identify the presence of lipid vacuoles (blue arrows). Note that *EphB2*^-/-^ Ad-MSC cultures contain more adipocytes and larger lipid vacuoles than WT or *EphB3*^-/-^ ones (200X, scale: 100 μm) (n=5, with 5 mice per group).

To confirm these adipogenic capacities, we cultured Ad-MSC in adipogenic medium for 21 days and stained the lipid vacuoles with *Oil Red* (**Fig. 3d**). *EphB2*^-/-^ Ad-MSC cultures contained more adipocytes with a larger number of lipid vacuoles than the remaining cells studied. Remarkably, *EphB3*^-/-^ Ad-MSC had a lower number of vacuoles than both WT and *EphB2*^-/-^ Ad-MSC.

For the analysis of osteogenic differentiation capacities, three pro-osteogenic transcription factors, *Runx2*, *Msx2* and *Osx*, were evaluated in the distinct Ad-MSC, cultured for 7 days in osteogenic medium. The expression of these factors significantly increased in *EphB3*^-/-^ Ad-MSC compared to that of both WT and *EphB2*^-/-^ values (**Fig. 4a-c**), highlighting the case of *Osx*, whose relative expression was 8-fold higher (**Fig. 4c**). On the contrary, the expression of these three factors did not vary in the *EphB2*^-/-^ Ad-MSC, compared with WT values (**Fig. 4a-c**).

**Fig. 4.**
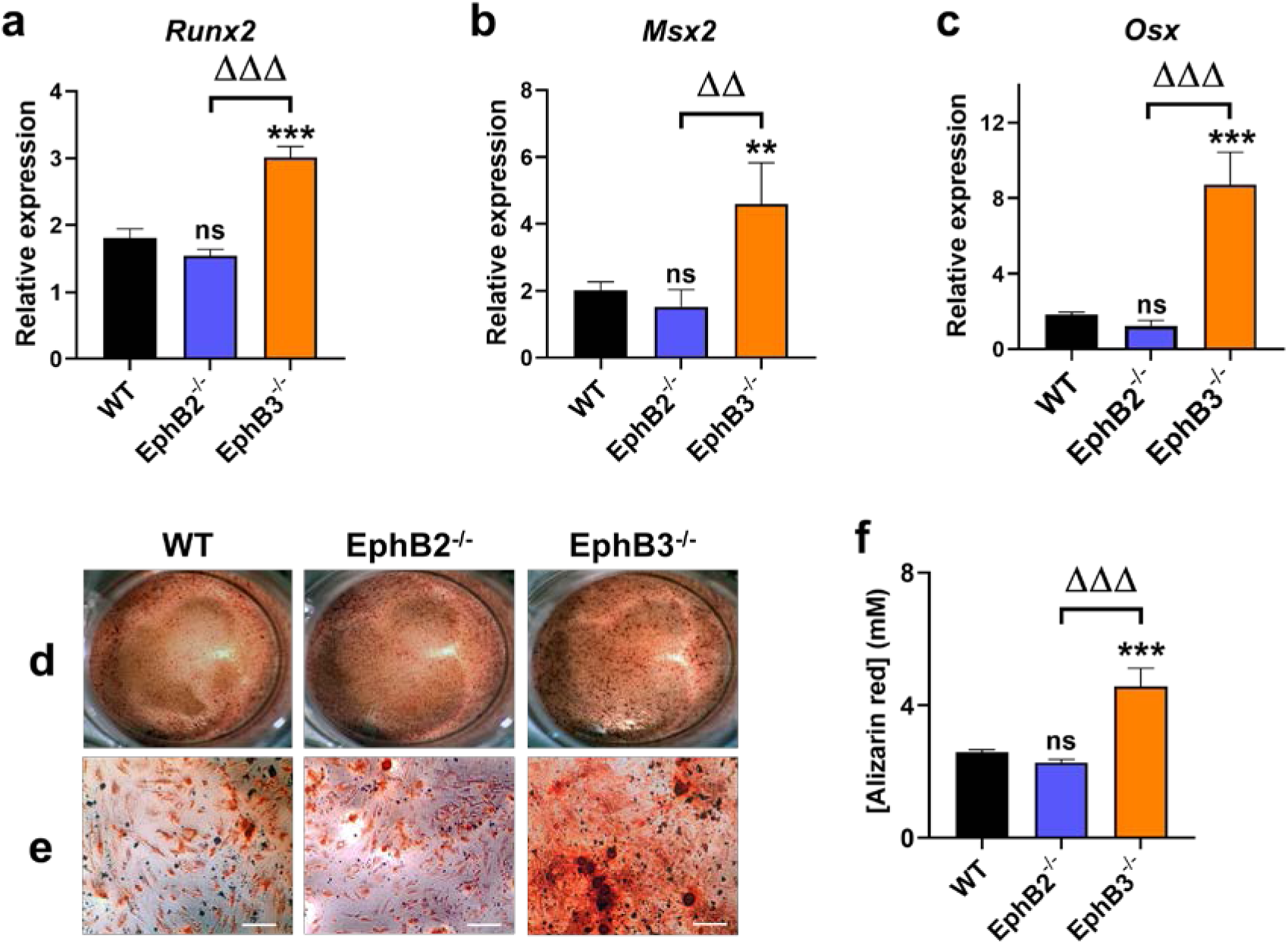
Osteogenic differentiation of both WT and *EphB*-deficient Ad-MSC. (a, b, c) Expression of osteogenic transcription factors in both differentiating WT and *EphB*-KO Ad-MSC. Note a significantly higher expression of the three osteogenic factors studied in the *EphB3*^-/-^ Ad-MSC as compared with WT and *EphB2*^-/-^ ones. (d, e) Evaluation of the osteogenic differentiation capacity by *Alizarin Red* staining after 21 days cultured with osteogenic medium (d. 2x; e. 100X, scale: 200 μm). (f) *Alizarin Red* quantification using CPC (562nm). *EphB3*^-/-^ Ad-MSC showed the highest osteogenic differentiation capacity. (ns=no significant difference, **/ΔΔ p value<0.01 and ***/ΔΔΔ p value<0.005) (n=5, with 5 mice per group).

In addition, the osteogenic differentiation capacity of both WT and *EphB*-KO Ad-MSC was analyzed using *Alizarin Red* to stain the calcium deposits formed after 21 days of culture. Thus, the results obtained by RT-qPCR were confirmed by a greater amount of calcium phosphate deposits being stained with *Alizarin Red* in the *EphB3*^-/-^ Ad-MSC than in the other Ad-MSC groups (**Fig. 4d, 4e**), as demonstrated with the staining quantification by CPC absorbance (**Fig. 4f**).

Altogether, these results confirmed that the *EphB3*^-/-^ Ad-MSC had a greater osteogenic potential than the other evaluated cells **(Fig. 4)**, while the *EphB2*^-/-^ ones had a higher potential for adipogenic differentiation **(Fig. 3)**.

### Altered gene expression of positive and negative osteogenic regulators in EphB3^-/-^ Ad-MSC

To explain the high osteogenic potential of *EphB3*-KO Ad-MSC, we analyzed the condition of several signaling pathways (i.e., BMP, Wnt/β*-*catenin and TGF-β) involved in the differentiation of MSC to bone cell lineage [6, 33]. After 7 days in osteogenic medium, we found an increased expression of several genes related to BMP signaling (*Bmp7*, *Bmpr1a*, *Bmpr1b* and *Smad9* (formerly called *Smad8*)) and no differences in others (*Bmpr2*, *Smad1*, *Smad5*, *Smad4* and *Smad6*) (**Fig. 5a, 5b**). Interestingly, the expression of some negative regulators of that pathway (*Bmp3*, *Smad7* and *Smurf1*) (**Fig. 5b**), as well as that of many BMP antagonists (*Chrd*, *Grem1*, *Grem2*, *Nog*, *Sost* and *Twsg*) (**Fig. 5c**) were, by contrast, significantly reduced in *EphB3*-KO Ad-MSC compared to WT. Apart from all these molecules, we also found an increase in *Wnt3a*, β-*catenin* and *Fst* transcripts (**Supplementary Fig. 1a-c**), but no significant differences in the expression of *Tgf*-β, *Erk* or *Bmp2*, and reduced values of *Bmp4* and *Bmp6 genes* (**Supplementary Fig. 1b, 1c**) compared with WT Ad-MSC values. To summarize, these results imply that the BMP7 signaling pathway, together with the decrease in several antagonists and negative regulators of this route, are key elements for the observed robust osteogenesis of the *EphB3*-KO Ad-MSC.

**Fig. 5.**
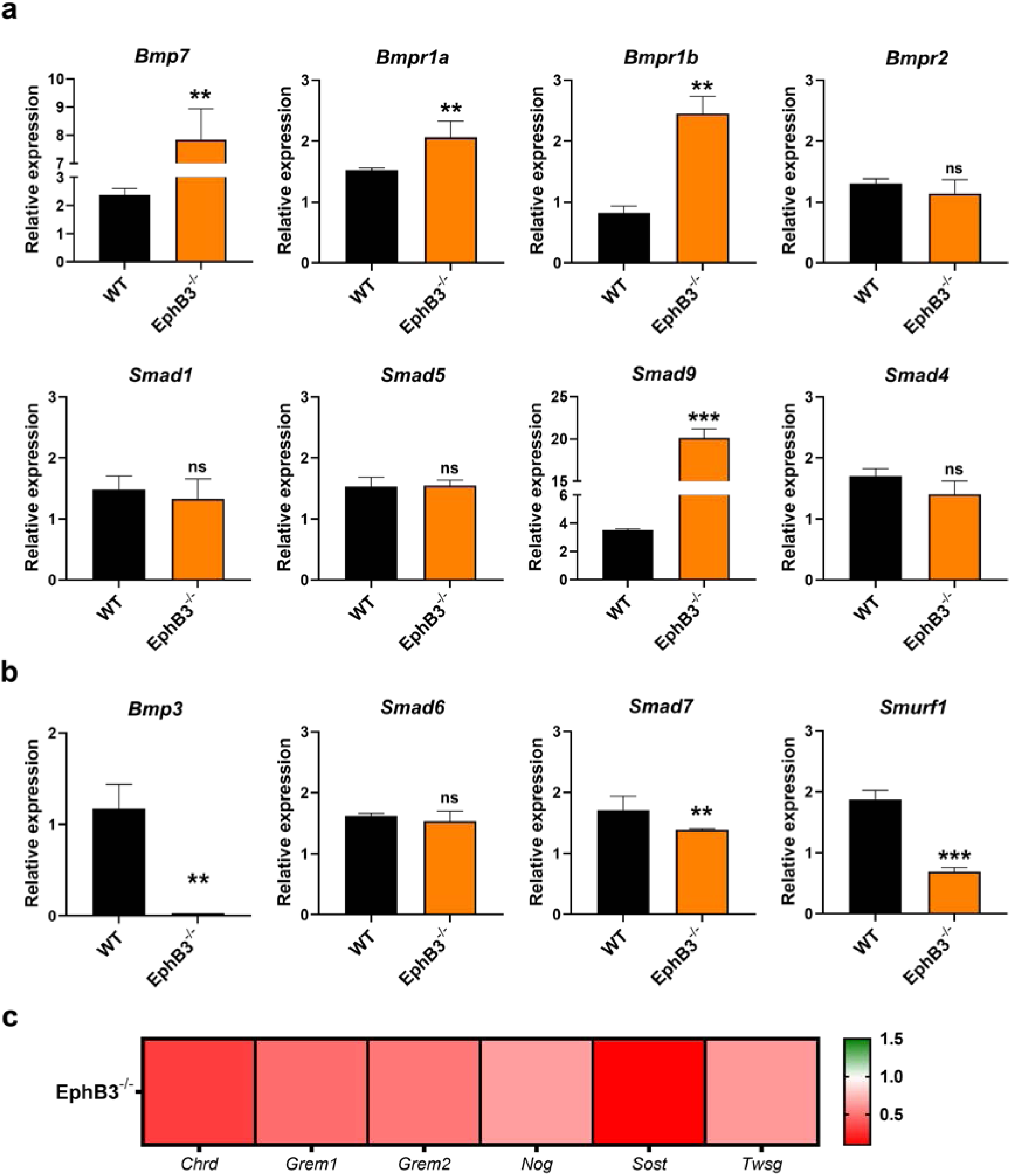
Gene expression analysis of BMP signaling pathway molecules in both WT and *EphB3*-KO Ad-MSC, 7 days after in vitro osteogenic differentiation. (a, b) Expression of BMP signaling route (a) and inhibitory molecules (b). (c) Heatmap of the expression of BMP antagonists, in which *EphB3*-KO and WT Ad-MSC data are compared (Chordin (*Chrd*), Gremlin (*Grem*), Noggin (*Nog*), Sclerostin (*Sost*) and Twisted gastrulation (*Twsg*)). Note the significantly increased expression of some BMP signaling effectors in *EphB3*-KO Ad-MSC, as well as an important decrease of both inhibitors and antagonists of the route, relative to WT expression (ns=no significant difference, ** p value<0.01 and *** p value<0.005) (n=3, with 5 mice per group).

### EphB3-deficient mice do not develop an osteoporotic phenotype

For a better understanding of the physiology of these mice, we analyzed their capacities to develop experimental osteoporosis. We chose two osteoporotic murine models (see Materials and Methods section), based on decreased estrogen levels by ovariectomy or long-term dexamethasone treatment. In the case of estrogen osteoporosis, the ovariectomized (OVX) condition was evaluated two months after surgery by measuring the serum estrogen concentration by an ELISA kit **(Supplementary Fig. 2)**. In the other experimental model, the osteoporotic condition was induced by injecting 2.5 mg/kg dexamethasone (DEX) daily for 14 days, being analyzed the day after.

Once mice had been sacrificed, femurs, second lumbar vertebras, urine and blood samples were collected for further analysis. One femur and the vertebra from each genotype were micro-CT scanned to quantify their condition as well as the bone volume fraction (BV/TV) and trabecular thickness (Tb.th). All untreated control mice (WT and *EphB* KO) showed no gross alterations in either bone morphology (**Fig. 6**) or other analyzed parameters (**Fig. 7**), with very similar values of BV/TV (over 8%) and Tb.th (around 0.12 mm). In the case of osteoporotic mice, WT and *EphB2*^-/-^ animals showed evident microarchitectural worsening of bone tissue (**Fig. 6**), according to the described phenotype for this pathology [34]. Remarkably, *EphB3*^-/-^ mice did not show any reminiscence of the typical loss of bone tissue in osteoporotic vertebrae (**Fig. 6**) or femur (**Supplementary Fig. 3**), with no morphological alterations in the micro-CT analysis, similarly to the condition of untreated mice. These observations were then confirmed by evaluating the BV/TV percentage and Tb.th value (**Fig. 7**). As expected, WT and *EphB2*^-/-^ treated mice showed a reduction in these parameters, compared to untreated control values, while *EphB3*^-/-^ mutants maintained significantly high values in the OVX and DEX groups. Interestingly, in the case of *EphB3*^-/-^ animals, these parameters were similar between untreated control and osteoporotic mutant mice, supporting the idea that the lack of EphB3 signaling protects from developing an osteoporotic condition, preventing the typical massive loss of bone tissue.

**Fig. 6.**
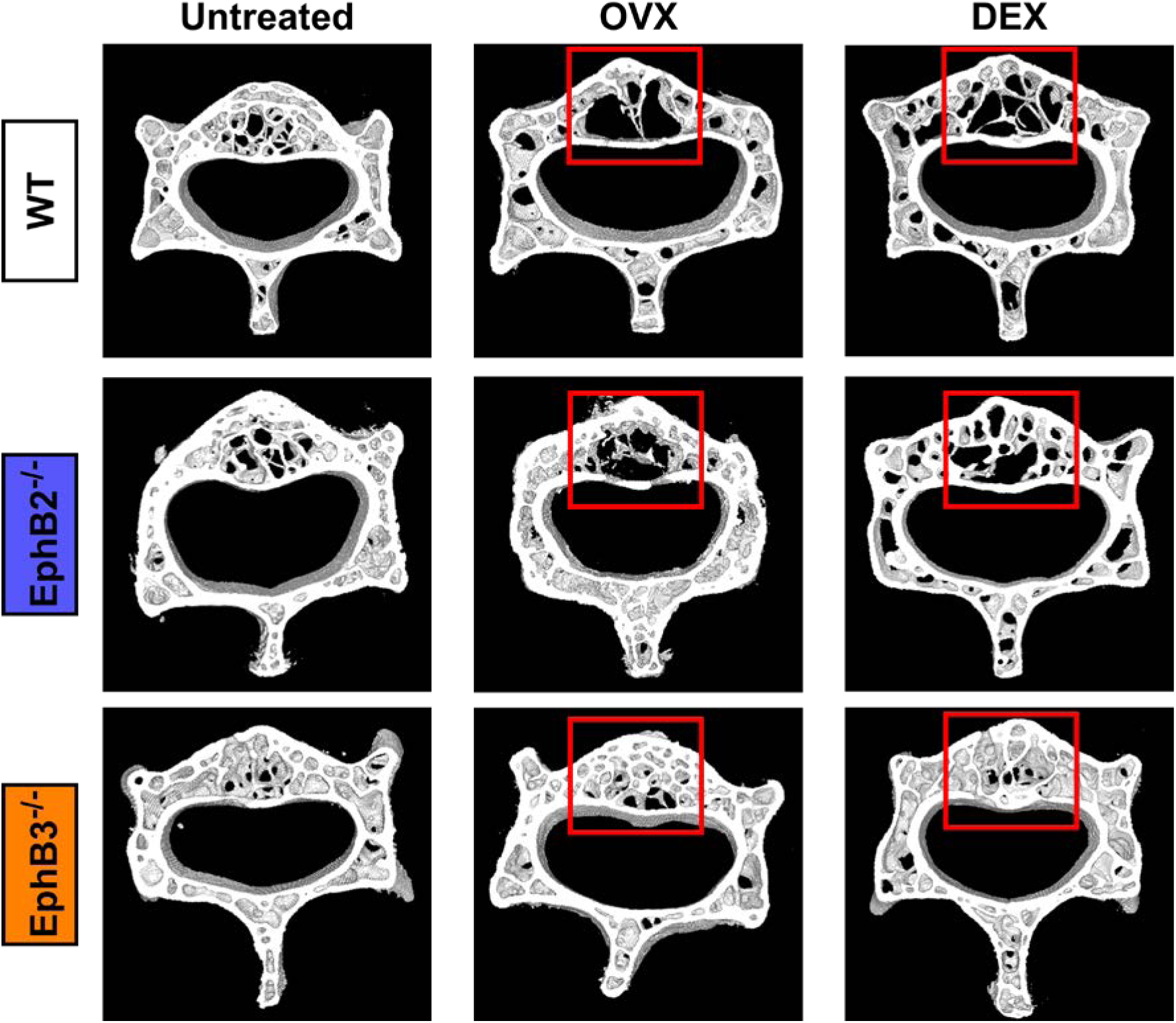
No bone alterations occur in *EphB3*^-/-^ mice after osteoporosis induction. Representative 3D reconstruction of de micro-CT images of vertebra scans. Showed analysis were made using the whole stack of the analyzed vertebras. All the treated WT and *EphB2*^-/-^ vertebras showed evident bone loss (see red squared areas), while the *EphB3*^-/-^ mice showed no bone alterations, like the untreated control mice (OVX: ovariectomized mice; DEX: dexamethasone treated mice) (n=6 mice per group).

**Fig. 7.**
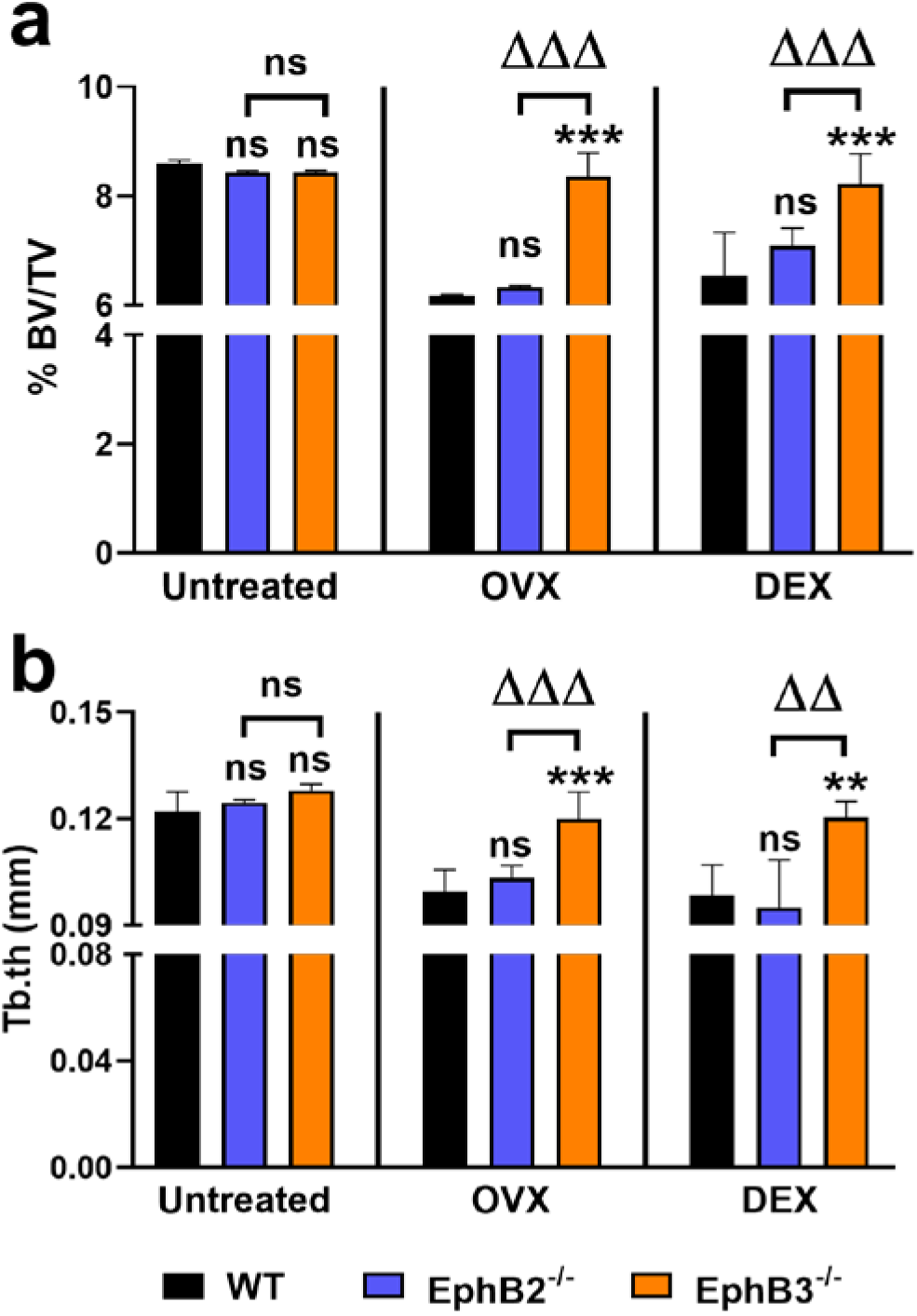
Induced-osteoporotic *EphB3*^-/-^ mice show similar bone morphometric values to untreated control animals. (a) Bone volume fraction (BV/TV) measured in vertebra micro-CTs. (b) Trabecular thickness (Tb.th) of vertebra micro-CTs. *EphB3*^-/-^ mice show significantly higher BV/TV and Tb.th values as compared with those of WT and *EphB2*^-/-^ mice, after OVX and DEX treatment, but have no differences when compared with the untreated control group (ns=no significant difference, **/ΔΔ p value<0.01 and ***/ΔΔΔ p value<0.005). For each condition statistical comparisons were made intra-and intergroups (OVX: ovariectomized mice; DEX: dexamethasone treated mice) (n=6 mice per group).

To confirm the existence of osteoporosis in the treated mice, we quantified the levels of Type I Collagen Cross-Linked C-Telopeptide (CTX-1) in serum of untreated control, OVX and DEX treated mice, as this peptide is a biomarker for bone resorption [35]. In both OVX and DEX treatment, the *EphB3*^-/-^ mice presented significantly lower values than osteoporotic WT and *EphB2*^-/-^ animals. In these two cases, as expected for this pathology, the serum CTX-1 concentration was higher than in the untreated controls (**Fig. 8a**). Together with CTX-1, we also quantified by ELISA the concentration of calcium (Ca^2+^) in urine (**Fig. 8b**), that tends to rise in osteoporosis. The Ca^2+^ concentration in urine also increased in all the ovariectomized mice, and in the ones treated with dexamethasone except for *EphB3*^-/-^ mice which presented significantly lower levels, like the untreated animals. Both results confirmed the lack of an osteoporotic phenotype in the *EphB3*^-/-^ treated mice.

**Fig. 8.**
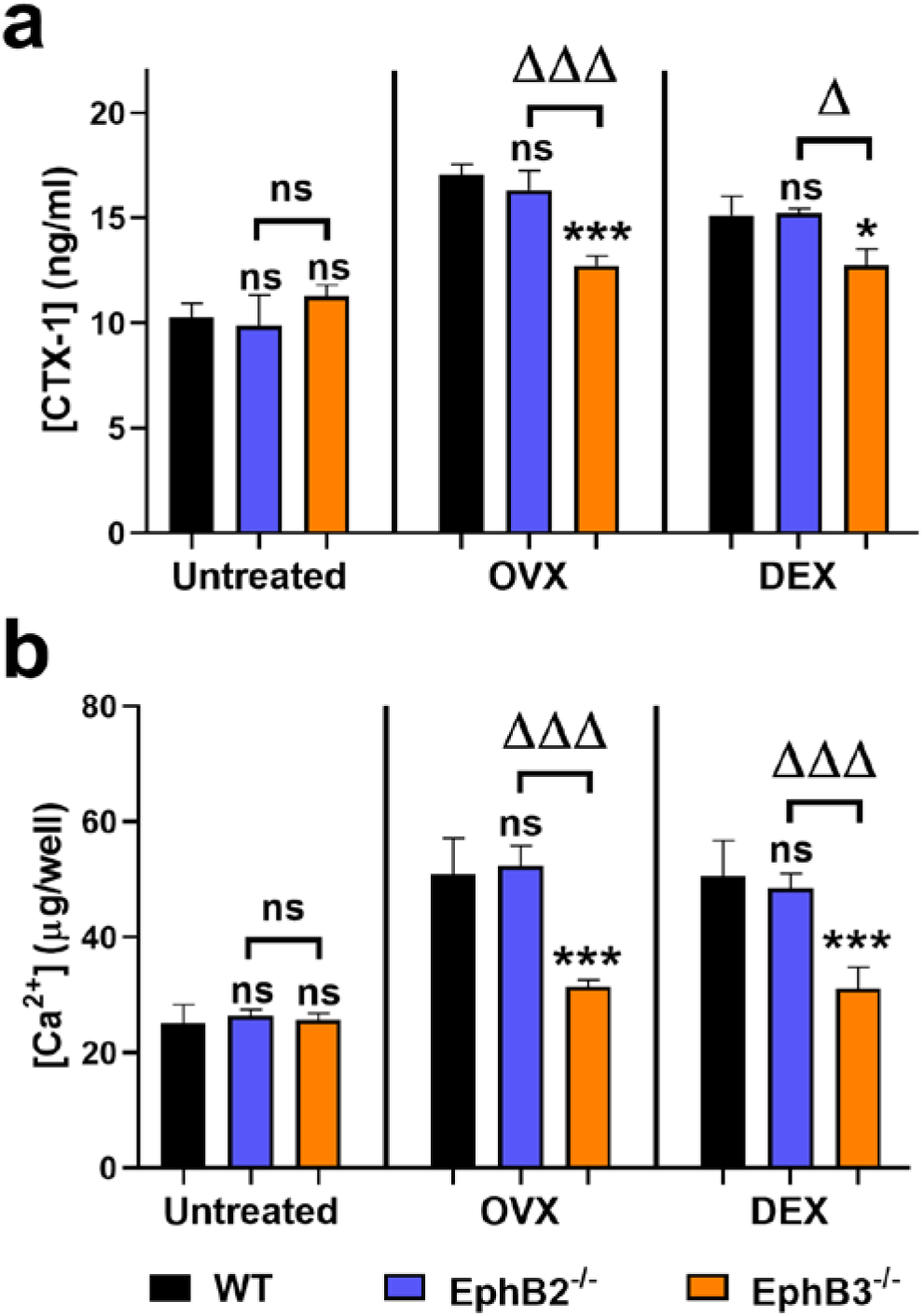
Decreased values of osteoporotic biomarkers in *EphB3*^-/-^ mice after OVX or DEX treatment. (a) CTX-1 levels in serum and (b) calcium concentration in urine of untreated control, OVX and DEX treated mice. In both osteoporotic groups, there is an increase in serum CTX-1 and urine calcium concentrations due to bone tissue degradation. In *EphB3*^-/-^ mice, the concentrations are lower than their counterparts, remaining unaltered as in the control condition. Values are compared statistically intra-and intergroups for each condition (ns=no significant difference, */Δ p value<0.05 and ***/ΔΔΔ p value<0.005) (OVX: ovariectomized mice; DEX: dexamethasone treated mice) (n=6 mice per group).

### Both increased proportions of osteoprogenitors and pre-osteoblasts, and reduced osteoclasts occur in EphB3-deficient mice which do not show osteoporotic features

The absence of an osteoporotic phenotype in *EphB3*^-/-^ mice could be due to any of these three non-exclusive reasons: their MSC differentiate and produce new bone more efficiently, there is a deficiency in the number or the activity of osteoclasts, or both. Accordingly, we studied by flow cytometry the proportions of these bone cell populations in the femur of untreated control and osteoporotic mice. As we previously found the same phenotype in OVX and DEX treated mice and applying the 3Rs Reduction principle (Directive 2010/63/EU), we decided to evaluate only the condition of DEX mice in these assays.

Our results showed that the percentages of both osteoprogenitor (Sca1^high^CD51^high^Lin^-^CD45^-^ CD31^-^) cells and pre-osteoblasts (Sca1^med^CD51^med^Lin^-^CD45^-^CD31^-^) were significantly higher in *EphB3*^-/-^ mice than in WT or *EphB2*^-/-^, in both untreated and osteoporotic mice (**Fig. 9a, 9b**). On the other hand, we also checked whether the osteoclast cell population was involved in the lack of an osteoporotic phenotype in *EphB3*^-/-^ mice. By flow cytometry, the percentage of these cells decreased in *EphB3*^-/-^ mice compared with WT and *EphB2*^-/-^, in both untreated and DEX conditions again (**Fig. 9c**). Next, to analyze their osteoclastogenic capacities, we used TRAP (Tartrate-Resistant Acid Phosphatase) staining to determine *in vitro* the number of osteoclasts after induced differentiation with RANKL, M-CSF and TNF-α (see detailed protocols in Materials and Methods section). In general terms, as expected, we found a significantly higher number of osteoclasts in the osteoporotic mice than in the control condition, as osteoclasts are enhanced by the glucocorticoid treatment. Notably, in both untreated and osteoporotic conditions, the number of osteoclasts in *EphB3*^-/-^ mice were lower than both WT and *EphB2*^-/-^ mice (**Fig. 9c**) and showed an impaired differentiation capacity after stimulation (**Fig. 9d**).

**Fig. 9.**
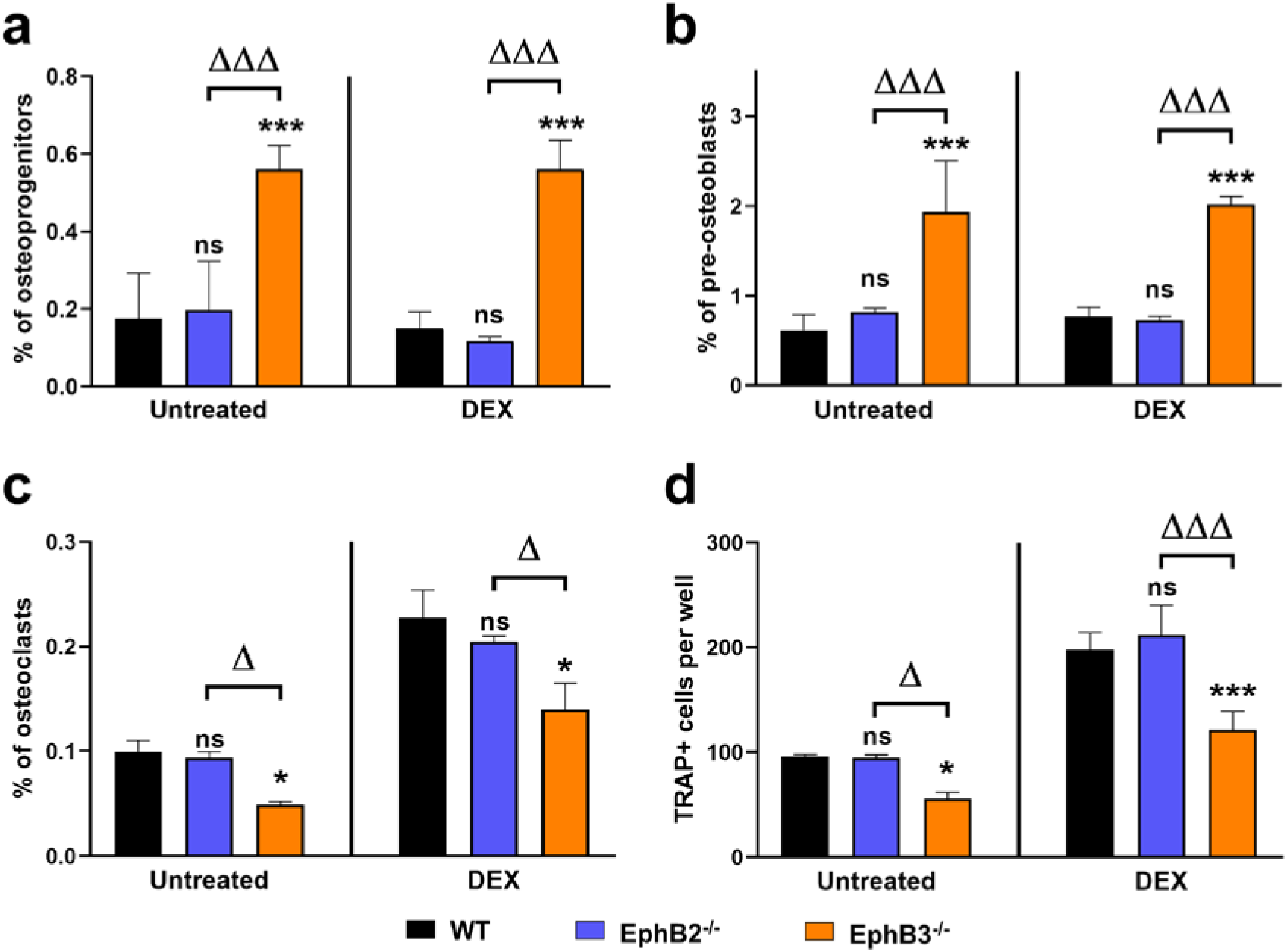
Femur bone cell populations analyzed by flow cytometry and TRAP staining. Percentage of (a) osteoprogenitors (Sca1^high^CD51^high^Lin^-^CD45^-^CD31^-^ cells), (b) pre-osteoblasts (Sca1^med^CD51^med^Lin^-^CD45^-^CD31^-^ cells) and (c) osteoclasts (cKit^+^CD115^+^CD3^-^CD45R^-^CD11b^-/low^), determined by flow cytometry. (d) Number of TRAP+ cells after 7 days of osteoclastic differentiation. The percentages of both osteoprogenitor cells (a) and pre-osteoblasts (b) were significantly higher in untreated and DEX *EphB3*^-/-^ mice compared with their respective WT. Also, the percentage of osteoclasts was significantly lower *in vi*vo (c), and after *in vitro* differentiation too (d), in *EphB3*^-/-^ mice compared with WT mice in untreated and DEX groups (ns=no significant difference, */Δ p value<0.05 and ***/ΔΔΔ p value<0.005). Statistical comparisons were made intra-and intergroups for each condition. DEX: dexamethasone-treated mice (n=5 mice per group).

In summary, EphB3 receptor could be involved in the suppression of osteoblast formation (osteoblastogenesis) and in promoting osteoclast differentiation (osteoclastogenesis). Accordingly, the lack of this signaling pathway in the *EphB3*^-/-^ mice coursed with a pro-osteogenic phenotype that was apparently sufficient to prevent experimentally-induced osteoporosis.

## Discussion

Osteogenesis is a complex process governed by multiple endogenous and external factors, including Eph tyrosine kinase receptors and their ligands, Ephrins [7]. This family of molecules plays a role in cell migration, adhesion and cellular attachment/detachment [16], which also contribute to embryonic patterning, skeletal development and bone homeostasis [7]. Unexpectedly, although the role played in these last processes by numerous Eph and Ephrins of both families A and B has been already studied, the relevance of EphB3 is scarcely known.

We [36] and other authors [9, 24] demonstrated the expression of most Eph and Ephrins A and B in MSC derived from bone marrow and adipose tissue. In the current study, we firstly examined the *in vitro* capacities of differentiation of Ad-MSC isolated from *EphB2* and *EphB3*-deficient mice compared with those of control, WT mice, revealing EphB3 receptor as an important suppressor of osteogenic differentiation.

The phenotypic characterization of both *EphB*-deficient Ad-MSC as well as their growth, cell cycle and apoptosis level, do not differ significantly from control, WT values, as we had evidenced previously [37], but the lack of EphB3 produces increased osteogenesis (higher expression of *Runx2*, *Msx2* and *Osterix* genes). On the contrary, *EphB2*^-/-^ Ad-MSC differentiate specially to adipose tissue and less efficiently than *EphB3*^-/-^ cells to bone lineage, but similarly to WT cells. In fact, EphB2 is claimed to be less important than other EphB for osteogenesis: *EphB2* global KO mice do not show significant skeletal alterations [18], and specific blocking peptides for EphB2 do not significantly reduce bone formation [24]. It is assumed that some pairs of Eph/Ephrins, particularly EphB4 and EphrinB2, are involved in bone homeostasis and regulate bidirectionally communication between osteoblasts and osteoclasts [7]. Thus, the activation of EphB4 signaling in osteoblasts promotes mineral formation [12], and blocking peptides impede its formation [24], while reverse signaling through its ligand EphrinB2 inhibits osteoclastogenesis [12]. *EphrinB1* also regulates osteogenesis, as it enhances bone formation after being overexpressed in osteoblast progenitors [38], it generates gross skeletal malformations in global and conditional knockout mice [18, 21, 39], and mice lacking *EphrinB1* in the osteogenic population develop an osteoporotic-like phenotype [40].

Regarding *EphB3*, apart from some seminal studies revealing its role in the organization of the secondary palate [25, 26], only recently, Kamath and Benson (2021) have reported its strong expression in adult calvaria sutures and suggested, in agreement with our current results, that EphB3 would be a limiting factor for osteogenesis [27]. Nevertheless, whereas Kamath and Benson [27] concluded that the observed changes in the skeleton of *EphB3* KO mice are minimal and presumably transient, our results conclusively demonstrate the remarkable capacity of *EphB3* KO mice for inducing a robust bone production.

In order to clarify how the absence of EphB3 signaling results in a high increase of the bone produced by Ad-MSC, we analyzed the expression of an extended group of genes known to be involved in the induction or blockade of bone cell differentiation [6]. Our analysis shows a dual effect of the lack of *EphB3* on Ad-MSC, consisting in an increased expression of some pro-osteogenic molecules of the BMP and Wnt/β-catenin pathways and reduced expression of numerous inhibitors of the BMP signaling pathway [6]. In the first group of molecules, the most notable variations were related to the increased expression of *Bmp7*. It is known that BMP7 induces the differentiation of MSC to bone cell lineage [41], and it has been claimed to be necessary to maintain osteogenesis [42], representing a potential target for fracture healing and osteoporosis treatment [43]. BMP7 expression is also reinforced by pro-osteogenic transcription factors such as Runx1 and Runx2 (the latter was also increased in *EphB3*^-/-^ Ad-MSC), establishing a positive feedback loop [44]. However, the expression of other pro-osteogenic factors, such as *Bmp4* and *Bmp6* was reduced. This could be correlated with the increase in *Bmp7*, as it has been demonstrated that the addition of BMP7 enhances osteogenic differentiation but inhibits endogenous *Bmp3*, *Bmp4* and *Bmp6* expressions in cultured BM-MSC [45]. This agrees with our findings in *EphB3*^-/-^ Ad-MSC where the expression of *Bmp3*, an inhibitor of BMP-mediated osteogenic signaling, is also totally downregulated [46].

BMP signaling is regulated by distinct mechanisms, including interference with either BMP ligands, BMP receptors or signal transducers of the Smad family [6]. Noggin (*Nog*) and Chordin (*Chrd*), two BMP antagonists that are downregulated in *EphB3*^-/-^ Ad-MSC, inhibit osteo-induction mediated by BMP2 and BMP4, but *Bmp7* expression is resistant to Nog inhibition [6]. The antagonistic effect of Sclerostin (Sost) on BMP signaling is controversial. The first studies supported a direct binding of Sost to BMP [47], but other studies showed its regulation to be mediated through the Wnt/BMP interaction [48]. Notably, *EphB3*^-/-^ Ad-MSC exhibited a high expression of *Follistatin* (*Fst*), another BMP antagonist. Indeed, this non-selective inhibitory molecule has recently been described, in agreement with our results, as an enhancer of osteogenic differentiation in Ad-MSC [49].

Finally, canonical BMP signaling is dependent on Smad intracellular transducers. Accordingly, we found a high increase in *Smad9* expression: a molecule that forms part of the Smad1/5/9 complex and, after interacting with Smad4, it translocates into the nucleus and activates the transcription of target genes, e.g. *Runx2* [6]. This major upregulation of *Smad9* expression, and to a lesser extent of *Smad1* and *Smad4*, after BMP7 stimulation, has been previously described by other authors in tenocyte-like cells [50]. In addition, *EphB3*^-/-^ Ad-MSC exhibited a downregulated expression of *Smad7* and *Smurf1*, two inhibitors of the BMP canonical pathway, whose presence impedes the correct formation of the Smad1/5/9-Smad4 complex [6]. These results, together with the absence of *Bmp3* expression, should favor enhanced *Bmp7* effects and, therefore, the higher osteogenic capacity observed in *EphB3*^-/-^ Ad-MSC.

Although there is no evidence for the relationship between EphB and BMP in the differentiation of MSC into bone cells, crosstalk between these two families of molecules has been reported in other tissues. The expression of EphB2, EphB3 and BMP antagonists in human colon crypts follows an opposite expression profile, compared to BMP positive regulators. Thus, in agreement with our current results, where the expression of EphB2 and EphB3 is low, BMP antagonists remain low as well, while the expression of BMP positive regulators is high [51]. Therefore, the pro-osteogenic phenotype of *EphB3*^-/-^ mice, summarized in **Fig. 10**, is a consequence of an altered balance of different positive and negative osteogenic signals that favors an augmented differentiation of Ad-MSC into the bone cell lineage.

**Fig. 10.**
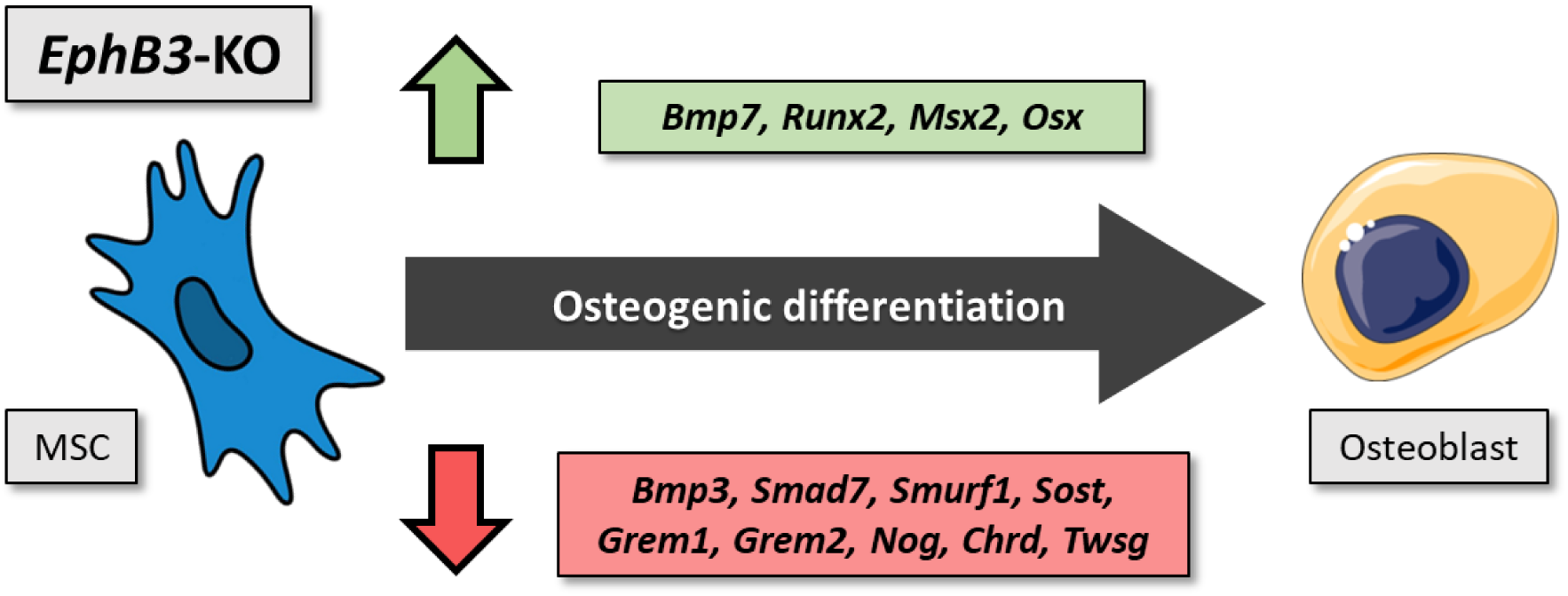
Gene expression changes related to the augmented osteogenic differentiation of *EphB3*-KO Ad-MSC. The lack of *EphB3* induces massive osteo-differentiation of MSC due to an increased expression of positive regulators (green box) and a notable reduction in antagonists (red box) of the BMP signaling pathway.

Apart from the signaling pathways involved in the *EphB3*^-/-^ Ad-MSC differentiation, we analyzed if the changes observed in the *EphB3*^-/-^ mice have some functional consequence. Remarkably, when we challenged the osteogenic process through two experimentally-induced osteoporotic models (ovariectomy or DEX treatment), we found that these *EphB3*-deficient mice do not develop any osteoporotic phenotype (bone loss and alterations in blood and urine parameters) and maintain high proportions of osteoprogenitors and low ones of osteoclasts, suggesting that the lack of EphB3 promotes overall bone formation. Thus, EphB3 would be an osteogenic repressor that is downregulated when bone formation is needed, as occurs in fracture bone healing [52], and is expressed in non-bone differentiating tissues, such as the proliferative chondrocyte zones [27].

Increased proportions of osteoprogenitors and pre-osteoblasts, together with reduced percentages of osteoclasts in both untreated control and treated *EphB3*^-/-^ mice, confirm that EphB3 presumably inhibits osteoblast differentiation from the early stages of maturation, as shown by our *in vitro* assays, whereas it supports osteoclastogenesis. Other Eph and Ephrins are also involved in these processes: EphB4 stimulates osteoblast differentiation whereas it inhibits osteoclastogenesis via EphrinB2-reverse signaling [12]. The role of Ephrins B in these processes has been repeatedly remarked. Both EphrinB1 [9] and EphrinB2 [53], the main ligands of EphB3 [54] are expressed in osteoblasts and osteoclasts [8] and have been described as negative regulators of osteoclast differentiation because their lack in osteoprogenitors results in increased numbers of osteoclasts [7]. In addition, Shimizu et al. analyzed the effects of alendronate, an anti-resorptive drug used in osteoporotic patients, affirming that it reduces osteoclast function and suppresses osteoblast differentiation, stimulating EphrinB1 expression in osteoclasts, together with that of EphB1 and EphB3 in osteoblasts. [55]. Furthermore,

EphrinB1-Fc stimulation of osteoblasts suppresses its differentiation, while EphrinB2-Fc enhances it, suggesting that interactions between EphrinB1 and EphB1/B3 may cause the inhibition of osteoblast differentiation [55].

According to our results, and under the premise demonstrated in many biological systems that the effects of Eph and Ephrin are a consequence of the global balance of signals transmitted by all the molecules expressed [16], we would like to propose a tentative, preliminary model for the role of Eph/EphrinB in murine osteogenesis **(Fig. 11).** Apart from the role of EphB3 in partially regulating first stages of MSC differentiation into bone cell lineage, as described above (**Fig. 10**), we think that in the presence of EphB2, EphB3 and EphB4 receptors, EphB3/EphrinB signaling negatively regulates the bone production induced by EphB4 and EphB2 forward signaling, with the concourse of reverse EphrinB1 and EphrinB2 signals. The lack of EphB2 in *EphB2*^-/-^ mice, might be compensated by EphB4, at least, showing the same osteogenic level as WT mice, both *in vitro* and *in vivo*. By contrast, in the absence of EphB3, bone production induced by the other two EphB would reach, if necessary, maximal levels without the “brake” of EphB3 signaling. Regarding the role of EphrinB1 and EphrinB2, they are, as indicated, the main ligands of the three described EphB, but their EphB-stimulation produces different results [55]. This same reasoning might be applied to the EphrinB signaling outcome when different EphB bind together: EphB4 and EphB2 interactions with EphrinB result in osteoclast inhibition [7], whereas EphB3 stimulation would enhance it. Indeed, to know the reason for these different Eph/Ephrin behaviors, it would be necessary to demonstrate that EphrinB1 and EphrinB2 follow different signaling pathways when stimulated by different EphB and *vice versa*.

**Fig. 11.**
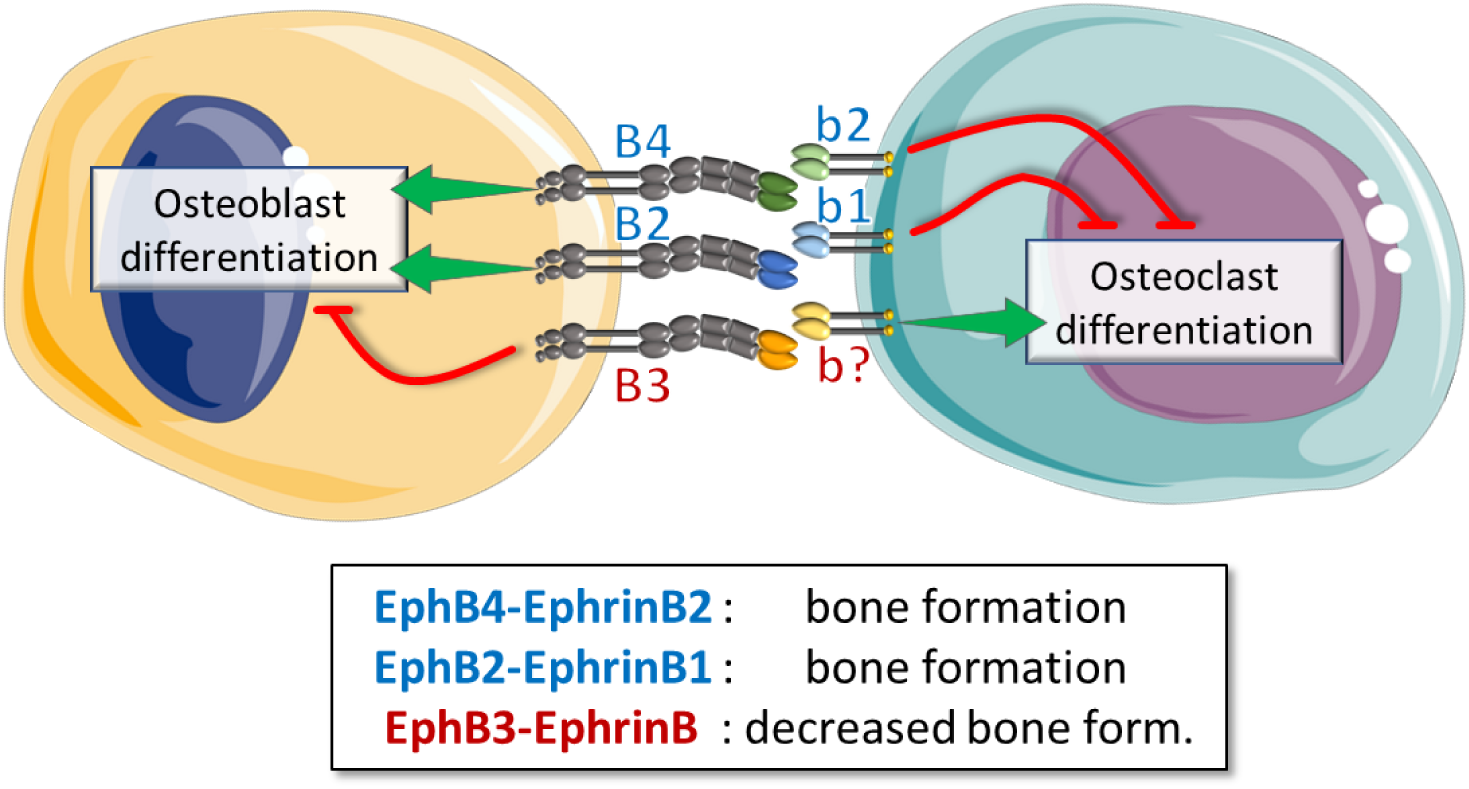
Integrative scheme on the role of Eph/Ephrins B in bone homeostasis. Forward signaling mediated by both EphB4 and EphB2 promotes (green arrows) osteoblast differentiation while the reverse signaling through their ligands, EphrinB2 and EphrinB1, inhibits (red lines) osteoclast development. On the other hand, EphB3 signaling would decrease osteoblast production (red line) whereas their ligands, EphrinB, would favor (green arrow) osteoclasts.

On the other hand, although our discussion mainly focuses on the Eph/Ephrin of family B, Eph/Ephrin of family A are also involved in bone homeostasis and development. Numerous members of this family are expressed in the bone cell lineage [7, 9, 23, 56], and reverse signaling through EphrinA2 has been described to enhance osteoclastogenesis whereas forward signal transmitted by EphA2 inhibits osteoblast differentiation [8, 12, 22]. Remarkably, available information suggests that the functions of Eph/Ephrin A in osteogenesis/bone resorption appear to be opposite to those generally assumed for Eph/Ephrin of family B. However, our current results support another view, as we propose herein for the EphB3/EphrinB pair.

In conclusion, our findings demonstrate the importance of EphB3 signaling in negatively regulating bone homeostasis. Thus, EphB3 would act early in the MSC differentiation process into bone progenitor cells, controlling the level of BMP signaling through their inhibitors and positive regulators (**Fig. 10**) and later, together with other EphB and Ephrins B, governing osteoblast maturation and osteoclastogenesis. Therefore, this receptor becomes a very important target to develop new therapies for bone pathologies and related diseases.

## Acknowledgments

We thank the Centers of Cytometry and Fluorescence Microscopy, Genomic and Animal Housing of the Complutense University of Madrid (UCM) for the use of their facilities.

## Statements & Declarations

### Author Contributions

Mariano R. Rodríguez-Sosa: experimental design, collection and assembly of data, data analyses and interpretation, and manuscript writing and reviewing. David Alfaro: conception of study, experimental design, collection and assembly of data, data analyses and interpretation, manuscript writing and reviewing. Luis M. del Castillo: collection and assembly of data, data analyses and interpretation. Adrián Belarra: micro-CT data collection. Agustín G. Zapata: funding acquisition, conception of study, data analyses and interpretation, manuscript writing and reviewing. All authors read and approved the final manuscript.

### Data Availability

The datasets used and/or analyzed during the current study are available from the corresponding author on reasonable request.

### Ethics approval

All procedures used in this study were approved by the Ethics Committee for Animal Research of the UCM and Regional Government of Madrid, following the official European guidelines for the care and use of laboratory animals (Directive 2010/63/UE) and the ARRIVE guidelines 2.0.

## Supplementary Information

**Supplementary Fig. 1.**
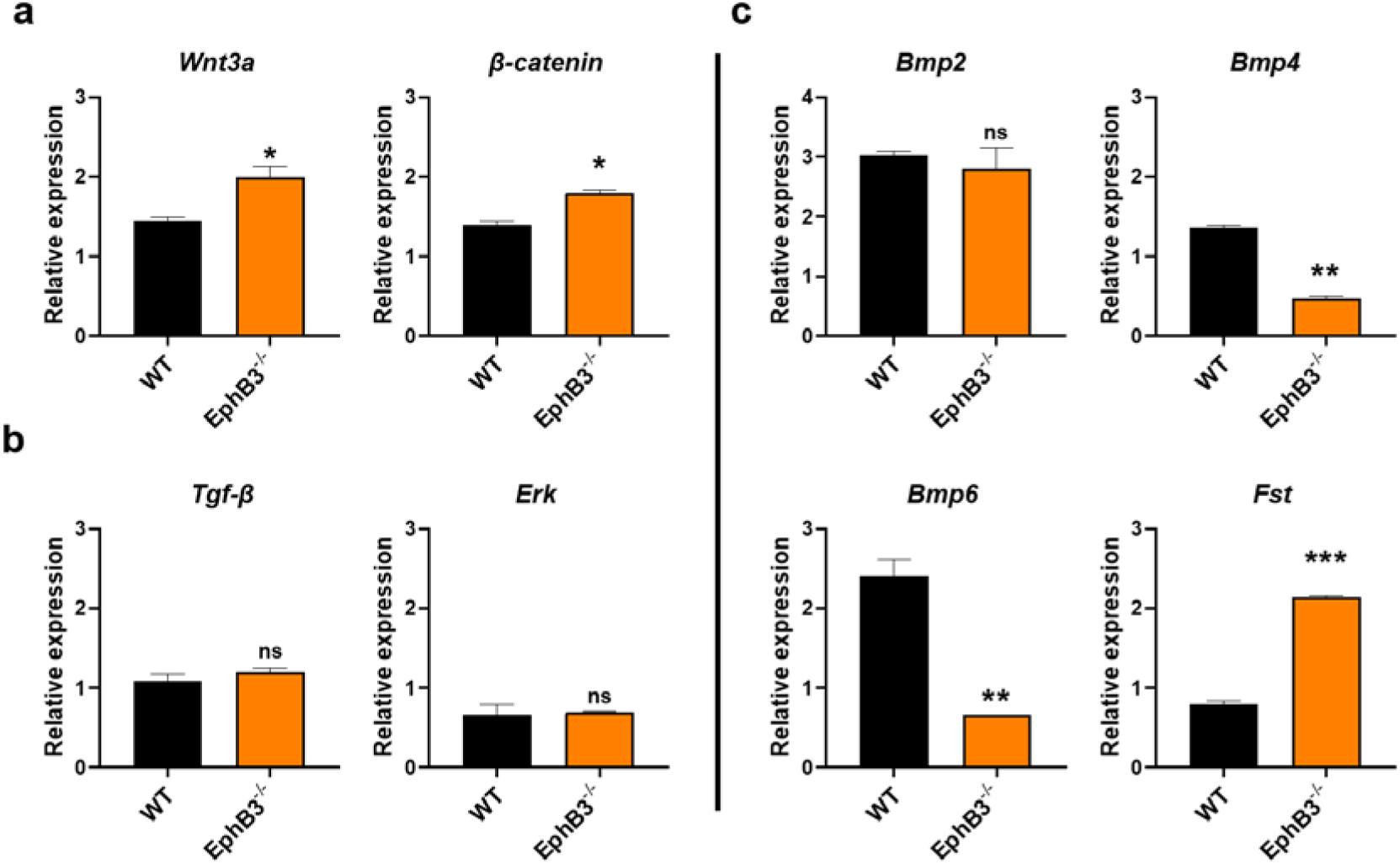
Title: Gene expression of different components of the Wnt/β-catenin, TGF-β and BMP signaling pathways in both WT and *EphB3*-KO Ad-MSC after 7 days in pro-osteogenic culture. **Description:** Whereas *Wnt3a*, β*-catenin* and *Follistatin* (*Fst*) transcripts increase significantly in *EphB3*-KO Ad-MSC, relative to WT ones (a), there are no variations in the expression of *Tgf-*β and *Erk* (b). Reduced relative expression of *Bmp4* and *Bmp6*, but not that of *Bmp2* (c) (* p value<0.05; ** p value<0.01) (n=3, with 5 mice per group).

**Supplementary Fig. 2.**
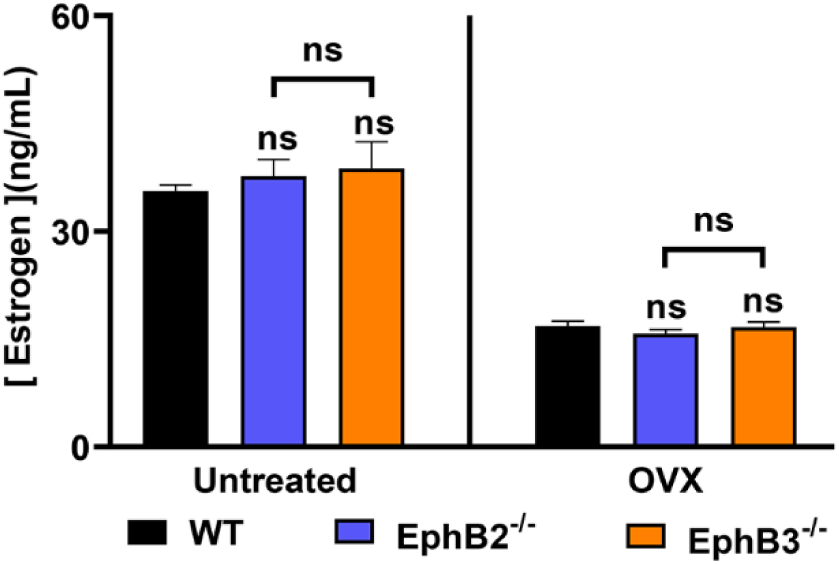
Title: Estrogen levels in the serum of control and OVX mice. **Description:** In all studied mice, the ovariectomy reduced the serum estrogen concentration. Intra-and intergroup statistical comparisons were performed for each condition (ns=no significant difference). OVX: ovariectomized mice. Note that significant differences occur between control and ovariectomized values but not between ovariectomized animals of different experimental groups (n=6 mice per group).

**Supplementary Fig. 3.**
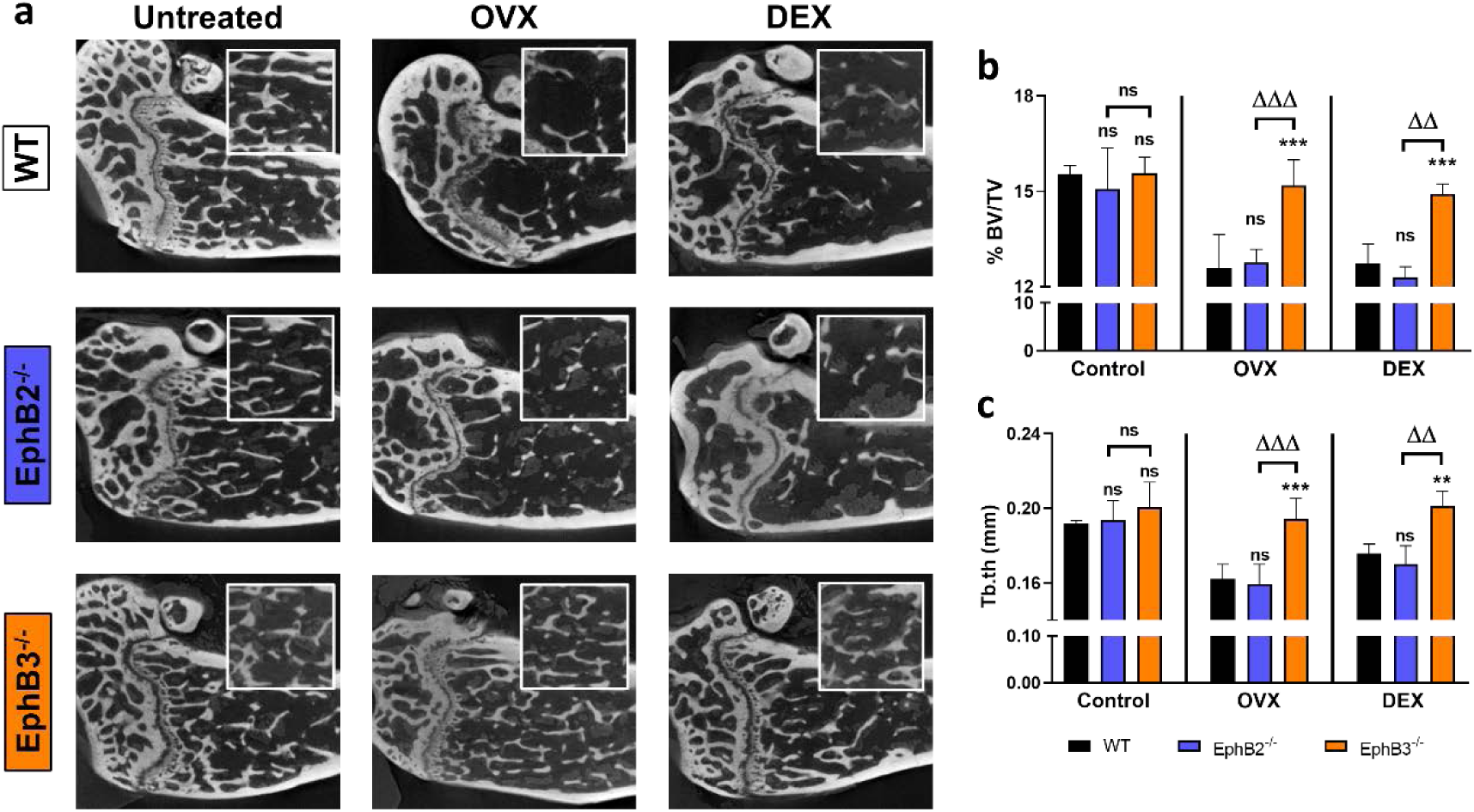
Title: No bone alterations occur in *EphB3*^-/-^ mice femurs after osteoporosis induction. **Description:** a. Representative images of micro-CT femur stacks. Quantification analyses were carried out by using the whole stack of the analyzed femurs. All the treated WT and *EphB2*^-/-^ femur showed evident bone loss, while the *EphB3*^-/-^ mice do not show bone alterations, as occurs in untreated, control mice. (b) Bone volume fraction (BV/TV) measured in femur micro-CTs. (c) Trabecular thickness (Tb.th) of femur micro-CTs. As in vertebra analyses, *EphB3*^-/-^ femurs show significantly higher BV/TV and Tb.th values than both WT and *EphB2*^-/-^ mice, after OVX and DEX, but have no differences when compared with the untreated control group (ns=no significant differences, **/ΔΔ p value<0.01 and ***/ΔΔΔ p value<0.005). For each condition, statistical intra-and intergroup (OVX: ovariectomized mice; DEX: dexamethasone treated mice) comparisons were carried out (n=6 mice per group).

## References

1. Sandona M, Di Pietro L, Esposito F, Ventura A, Silini AR, Parolini O, Saccone V: Mesenchymal Stromal Cells and Their Secretome: New Therapeutic Perspectives for Skeletal Muscle Regeneration. Front Bioeng Biotechnol 2021, 9:652970.

2. Dominici MLB, K.; Mueller, I.; Slaper-Cortenbach, I.; Marini, F.; Krause, D.; Deans, R.; Keating, A.; Prockop, D.; Horwitz, E.: Minimal criteria for defining multipotent mesenchymal stromal cells. The International Society for Cellular Therapy position statement. Cytotherapy 2006, 8:315–317.

3. Garcia-Bernal D, Garcia-Arranz M, Yanez RM, Hervas-Salcedo R, Cortes A, Fernandez-Garcia M, Hernando-Rodriguez M, Quintana-Bustamante O, Bueren JA, Garcia-Olmo D et al: The Current Status of Mesenchymal Stromal Cells: Controversies, Unresolved Issues and Some Promising Solutions to Improve Their Therapeutic Efficacy. Front Cell Dev Biol 2021, 9:650664.

4. Krampera M, Le Blanc K: Mesenchymal stromal cells: Putative microenvironmental modulators become cell therapy. Cell Stem Cell 2021, 28(10):1708–1725.

5. Soliman H, Theret M, Scott W, Hill L, Underhill TM, Hinz B, Rossi FMV: Multipotent stromal cells: One name, multiple identities. Cell Stem Cell 2021, 28(10):1690–1707.

6. Thomas S, Jaganathan BG: Signaling network regulating osteogenesis in mesenchymal stem cells. J Cell Commun Signal 2022, 16(1):47–61.

7. Arthur A, Gronthos S: Eph-Ephrin Signaling Mediates Cross-Talk Within the Bone Microenvironment. Front Cell Dev Biol 2021, 9:598612.

8. Matsuo K, Otaki N: Bone cell interactions through Eph/ephrin: bone modeling, remodeling and associated diseases. Cell Adh Migr 2012, 6(2):148–156.

9. Nguyen TM, Arthur A, Gronthos S: The role of Eph/ephrin molecules in stromal-hematopoietic interactions. Int J Hematol 2016, 103(2):145–154.

10. Salvucci O, Maric D, Economopoulou M, Sakakibara S, Merlin S, Follenzi A, Tosato G: EphrinB reverse signaling contributes to endothelial and mural cell assembly into vascular structures. Blood 2009, 114(8):1707–1716.

11. Tonna S, Sims NA: Talking among ourselves: paracrine control of bone formation within the osteoblast lineage. Calcif Tissue Int 2014, 94(1):35–45.

12. Zhao C, Irie N, Takada Y, Shimoda K, Miyamoto T, Nishiwaki T, Suda T, Matsuo K: Bidirectional ephrinB2-EphB4 signaling controls bone homeostasis. Cell Metab 2006, 4(2):111–121.

13. Hadjimichael AC, Pergaris A, Kaspiris A, Foukas AF, Kokkali S, Tsourouflis G, Theocharis S: The EPH/Ephrin System in Bone and Soft Tissue Sarcomas’ Pathogenesis and Therapy: New Advancements and a Literature Review. Int J Mol Sci 2022, 23(9).

14. Taylor H, Campbell J, Nobes CD: Ephs and ephrins. Curr Biol 2017, 27(3):R90–R95.

15. Darling TK, Lamb TJ: Emerging Roles for Eph Receptors and Ephrin Ligands in Immunity. Front Immunol 2019, 10:1473.

16. Kania A, Klein R: Mechanisms of ephrin-Eph signalling in development, physiology and disease. Nat Rev Mol Cell Biol 2016, 17(4):240–256.

17. Niethamer TK, Bush JO: Getting direction(s): The Eph/ephrin signaling system in cell positioning. Dev Biol 2019, 447(1):42–57.

18. Compagni A, Logan M, Klein R, Adams RH: Control of skeletal patterning by ephrinB1-EphB interactions. Dev Cell 2003, 5(2):217–230.

19. Davy A, Bush JO, Soriano P: Inhibition of gap junction communication at ectopic Eph/ephrin boundaries underlies craniofrontonasal syndrome. PLoS Biol 2006, 4(10):e315.

20. Twigg SR, Matsumoto K, Kidd AM, Goriely A, Taylor IB, Fisher RB, Hoogeboom AJ, Mathijssen IM, Lourenco MT, Morton JE et al: The origin of EFNB1 mutations in craniofrontonasal syndrome: frequent somatic mosaicism and explanation of the paucity of carrier males. Am J Hum Genet 2006, 78(6):999–1010.

21. Xing W, Kim J, Wergedal J, Chen ST, Mohan S: Ephrin B1 regulates bone marrow stromal cell differentiation and bone formation by influencing TAZ transactivation via complex formation with NHERF1. Mol Cell Biol 2010, 30(3):711–721.

22. Irie N, Takada Y, Watanabe Y, Matsuzaki Y, Naruse C, Asano M, Iwakura Y, Suda T, Matsuo K: Bidirectional signaling through ephrinA2-EphA2 enhances osteoclastogenesis and suppresses osteoblastogenesis. J Biol Chem 2009, 284(21):14637–14644.

23. Yamada T, Yuasa M, Masaoka T, Taniyama T, Maehara H, Torigoe I, Yoshii T, Shinomiya K, Okawa A, Sotome S: After repeated division, bone marrow stromal cells express inhibitory factors with osteogenic capabilities, and EphA5 is a primary candidate. Bone 2013, 57(2):343–354.

24. Arthur A, Zannettino A, Panagopoulos R, Koblar SA, Sims NA, Stylianou C, Matsuo K, Gronthos S: EphB/ephrin-B interactions mediate human MSC attachment, migration and osteochondral differentiation. Bone 2011, 48(3):533–542.

25. Orioli D, Henkemeyer M, Lemke G, Klein R, Pawson T: Sek4 and Nuk receptors cooperate in guidance of commissural axons and in palate formation. EMBO J 1996, 15(22):6035–6049.

26. Risley M, Garrod D, Henkemeyer M, McLean W: EphB2 and EphB3 forward signalling are required for palate development. Mech Dev 2009, 126(3-4):230–239.

27. Kamath RAD, Benson MD: EphB3 as a potential mediator of developmental and reparative osteogenesis. Cells Tissues Organs 2021.

28. Sozen T, Ozisik L, Basaran NC: An overview and management of osteoporosis. Eur J Rheumatol 2017, 4(1):46–56.

29. Gruber HE, Somayaji S, Riley F, Hoelscher GL, Norton HJ, Ingram J, Hanley EN, Jr.: Human adipose-derived mesenchymal stem cells: serial passaging, doubling time and cell senescence. Biotech Histochem 2012, 87(4):303–311.

30. Baryawno N, Przybylski D, Kowalczyk MS, Kfoury Y, Severe N, Gustafsson K, Kokkaliaris KD, Mercier F, Tabaka M, Hofree M et al: A Cellular Taxonomy of the Bone Marrow Stroma in Homeostasis and Leukemia. Cell 2019, 177(7):1915–1932 e1916.

31. Tevlin R, McArdle A, Chan CK, Pluvinage J, Walmsley GG, Wearda T, Marecic O, Hu MS, Paik KJ, Senarath-Yapa K et al: Osteoclast derivation from mouse bone marrow. J Vis Exp 2014(93):e52056.

32. Domander R, Felder AA, Doube M: BoneJ2 - refactoring established research software. Wellcome Open Res 2021, 6:37.

33. Zakrzewski W, Dobrzynski M, Szymonowicz M, Rybak Z: Stem cells: past, present, and future. Stem Cell Res Ther 2019, 10(1):68.

34. Akbar MA, Lu Y, Elshikha AS, Chen MJ, Yuan Y, Whitley EM, Holliday LS, Chang LJ, Song S: Transplantation of Adipose Tissue-Derived Mesenchymal Stem Cell (ATMSC) Expressing Alpha-1 Antitrypsin Reduces Bone Loss in Ovariectomized Osteoporosis Mice. Hum Gene Ther 2017, 28(2):179–189.

35. Ersek A, Santo AI, Vattakuzhi Y, George S, Clark AR, Horwood NJ: Strain dependent differences in glucocorticoid-induced bone loss between C57BL/6J and CD-1 mice. Sci Rep 2016, 6:36513.

36. Alfaro D, Zapata AG: Eph/Ephrin-mediated stimulation of human bone marrow mesenchymal stromal cells correlates with changes in cell adherence and increased cell death. Stem Cell Res Ther 2018, 9(1):172.

37. Alfaro D, Rodriguez-Sosa MR, Zapata AG: Eph/ephrin Signaling and Biology of Mesenchymal Stromal/Stem Cells. J Clin Med 2020, 9(2).

38. Cheng S, Kesavan C, Mohan S, Qin X, Alarcon CM, Wergedal J, Xing W: Transgenic overexpression of ephrin b1 in bone cells promotes bone formation and an anabolic response to mechanical loading in mice. PLoS One 2013, 8(7):e69051.

39. Nguyen TM, Arthur A, Paton S, Hemming S, Panagopoulos R, Codrington J, Walkley CR, Zannettino AC, Gronthos S: Loss of ephrinB1 in osteogenic progenitor cells impedes endochondral ossification and compromises bone strength integrity during skeletal development. Bone 2016, 93:12–21.

40. Arthur A, Nguyen TM, Paton S, Klisuric A, Zannettino ACW, Gronthos S: The osteoprogenitor-specific loss of ephrinB1 results in an osteoporotic phenotype affecting the balance between bone formation and resorption. Sci Rep 2018, 8(1):12756.

41. Chen G, Deng C, Li YP: TGF-beta and BMP signaling in osteoblast differentiation and bone formation. Int J Biol Sci 2012, 8(2):272–288.

42. Shea CM, Edgar CM, Einhorn TA, Gerstenfeld LC: BMP treatment of C3H10T1/2 mesenchymal stem cells induces both chondrogenesis and osteogenesis. J Cell Biochem 2003, 90(6):1112–1127.

43. Reddi AH: Bone morphogenetic proteins: from basic science to clinical applications. J Bone Joint Surg Am 2001, 83-A Suppl 1(Pt 1):S1-6.

44. Tang CY, Wu M, Zhao D, Edwards D, McVicar A, Luo Y, Zhu G, Wang Y, Zhou HD, Chen W et al: Runx1 is a central regulator of osteogenesis for bone homeostasis by orchestrating BMP and WNT signaling pathways. PLoS Genet 2021, 17(1):e1009233.

45. Edgar CM, Chakravarthy V, Barnes G, Kakar S, Gerstenfeld LC, Einhorn TA: Autogenous regulation of a network of bone morphogenetic proteins (BMPs) mediates the osteogenic differentiation in murine marrow stromal cells. Bone 2007, 40(5):1389–1398.

46. Aluganti Narasimhulu C, Singla DK: The Role of Bone Morphogenetic Protein 7 (BMP-7) in Inflammation in Heart Diseases. Cells 2020, 9(2).

47. Winkler DG, Sutherland MK, Geoghegan JC, Yu C, Hayes T, Skonier JE, Shpektor D, Jonas M, Kovacevich BR, Staehling-Hampton K et al: Osteocyte control of bone formation via sclerostin, a novel BMP antagonist. EMBO J 2003, 22(23):6267–6276.

48. Kamiya N, Ye L, Kobayashi T, Mochida Y, Yamauchi M, Kronenberg HM, Feng JQ, Mishina Y: BMP signaling negatively regulates bone mass through sclerostin by inhibiting the canonical Wnt pathway. Development 2008, 135(22):3801–3811.

49. Wallner C, Huber J, Drysch M, Schmidt SV, Wagner JM, Dadras M, Dittfeld S, Becerikli M, Jaurich H, Lehnhardt M et al: Activin Receptor 2 Antagonization Impairs Adipogenic and Enhances Osteogenic Differentiation in Mouse Adipose-Derived Stem Cells and Mouse Bone Marrow-Derived Stem Cells In Vitro and In Vivo. Stem Cells Dev 2019, 28(6):384–397.

50. Klatte-Schulz F, Giese G, Differ C, Minkwitz S, Ruschke K, Puts R, Knaus P, Wildemann B: An investigation of BMP-7 mediated alterations to BMP signalling components in human tenocyte-like cells. Sci Rep 2016, 6:29703.

51. Kosinski C, Li VS, Chan AS, Zhang J, Ho C, Tsui WY, Chan TL, Mifflin RC, Powell DW, Yuen ST et al: Gene expression patterns of human colon tops and basal crypts and BMP antagonists as intestinal stem cell niche factors. Proc Natl Acad Sci U S A 2007, 104(39):15418–15423.

52. Kaur A, Xing W, Mohan S, Rundle CH: Changes in ephrin gene expression during bone healing identify a restricted repertoire of ephrins mediating fracture repair. Histochem Cell Biol 2019, 151(1):43–55.

53. Tonna S, Poulton IJ, Taykar F, Ho PW, Tonkin B, Crimeen-Irwin B, Tatarczuch L, McGregor NE, Mackie EJ, Martin TJ et al: Chondrocytic ephrin B2 promotes cartilage destruction by osteoclasts in endochondral ossification. Development 2016, 143(4):648–657.

54. Noberini R, Rubio de la Torre E, Pasquale EB: Profiling Eph receptor expression in cells and tissues: a targeted mass spectrometry approach. Cell Adh Migr 2012, 6(2):102–112.

55. Shimizu E, Tamasi J, Partridge NC: Alendronate affects osteoblast functions by crosstalk through EphrinB1-EphB. J Dent Res 2012, 91(3):268–274.

56. Yamada T, Yoshii T, Yasuda H, Okawa A, Sotome S: Dexamethasone Regulates EphA5, a Potential Inhibitory Factor with Osteogenic Capability of Human Bone Marrow Stromal Cells. Stem Cells Int 2016, 2016:1301608.

